# Gram-positive and Gram-negative Bacteria Share Common Principles to Coordinate Growth and the Cell Cycle at the Single-cell Level

**DOI:** 10.1101/726596

**Authors:** John T. Sauls, Sarah E. Cox, Quynh Do, Victoria Castillo, Zulfar Ghulam-Jelani, Suckjoon Jun

## Abstract

*Bacillus subtilis* and *Escherichia coli* are evolutionarily divergent model organisms that have elucidated fundamental differences between Gram-positive and Gram-negative bacteria, respectively. Despite their differences in cell cycle control at the molecular level, both organisms follow the same phenomenological principle for cell size homeostasis known as the adder. We thus asked to what extent *B. subtilis* and *E. coli* share common physiological principles in coordinating growth and the cell cycle. To answer this question, we measured physiological parameters of *B. subtilis* under various steady-state growth conditions with and without translation inhibition at both population and single-cell level. These experiments revealed core shared physiological principles between *B. subtilis* and *E. coli*. Specifically, we show that both organisms maintain an invariant cell size per replication origin at initiation, with and without growth inhibition, and even during nutrient shifts at the single-cell level. Furthermore, both organisms also inherit the same “hierarchy” of physiological parameters ranked by their coefficient of variation. Based on these findings, we suggest that the basic coordination principles between growth and the cell cycle in bacteria may have been established in the very early stages of evolution.

## Introduction

Our current understanding of fundamental, quantitative principles in bacterial physiology is largely based on studies of *Escherichia coli* and *Salmonella typhimurium* (see Jun et al.^1^ for a review of the history and recent progress). Several of these principles have been presented in the form of ‘growth laws’^1^. For example, the ‘growth law of cell size’ states that the average cell size increases exponentially with respect to the nutrient-imposed growth rate^2^. This principle has been extended to the ‘growth law of unit cells’, which allows prediction of the average cell size based on the growth rate and the cell cycle duration for any steady-state growth condition^3^.

Gram-positive *B. subtilis* is distinct from Gram-negative *E. coli* at the genetic, molecular, and regulatory level^4^. However, despite their evolutionary divergence, *B. subtilis* and *E. coli* follow the same phenomenological principle of cell-size homeostasis known as the adder principle^5, 6^. Furthermore, both organisms share the identical mechanistic origin of the adder principle, namely, a molecular threshold for division proteins and their balanced biosynthesis during growth^7^. Based on these findings, we wanted to know to what extent *B. subtilis* and *E. coli* coordinate growth, size, and cell cycle in the same manner. A shared coordination framework would imply that, despite phylogenetic and molecular diversity, physiological regulation in bacteria is functionally conserved.

In order to create a full complement of data necessary for comparative analysis, we measured the growth and cell cycle parameters of *B. subtilis* at both the population and single-cell level under a wide range of conditions.

Previous population-level studies have found that *B. subtilis*, like *E. coli*, initiates replication at a fixed mass, establishing a regulatory bridge between cell size and cell cycle control^8–10^. We extended this avenue with single-cell methods to precisely measure the cell cycle parameters in individual *B. subtilis* cells across conditions^7, 11^. These results showed that the initiation size is constant not only in steady-state conditions, but also during nutrient shifts between two steady-state conditions. This strongly supports a threshold model for initiation in both steady and dynamic environments^3, 7, 12, 13^.

The single-cell approach also allowed us to compare the relative variability of all growth and cell cycle parameters both between conditions and between species. These measurements reveal a strikingly similar hierarchy of physiological parameters between *B. subtilis* and *E. coli* in terms of tightness of their control.

The richness of our quantitative physiological data generated in *B. subtilis* is comparable to that in *E. coli*, providing key evidence that *B. subtilis* and *E. coli* share core phenomenological and quantitative principles that govern their physiology, thereby providing a unified picture of bacterial growth, size, and cell cycle coordination.

## Results and discussion

### Ensuring steady-state growth in *B. subtilis*

Maintaining a steady-state growth is essential for reproducible measurements of the physiological state of the cell^1^. In steady-state growth, the total biomass of the culture increases exponentially with time and protein biosynthesis is balanced with the total biomass increase. That is, the protein production rate is the same as the growth rate of the cell. As a result, average protein concentrations are constant, whereas the total amount of proteins increases in proportion to the cell volume. This constant concentration and proportional increase also applies to other macromolecules such as DNA, RNA, phospholipids, and the cell wall.

To achieve steady-state measurements in *B. subtilis*, we grew and monitored cells over many generations using a multiplex turbidostat that we previously used for *E. coli*^3^ (Figure **1**A). For both population and single-cell methods, we began cultures from single colonies and pre-cultured cells using appropriate batch methods before transferring to continuous culture set-ups (Materials and methods). We ensured pre-cultures did not enter stationary phase to avoid sporulation. We used a *B. subtilis* strain which was non-motile and non-biofilm forming to facilitate single cell-size measurements. This was necessary because *B. subtilis* exhibits a temporal chaining phenotype, particularly in faster growth conditions^14, 15^. During chaining, cells are physically connected yet their cytoplasms are compartmentalized, obfuscating a definition of division^16, 17^. Our strain contained a genetic modification to abolish cell chaining, ensuring that cell separation coincided with septation^18^ (Materials and methods).

**Figure 1:**
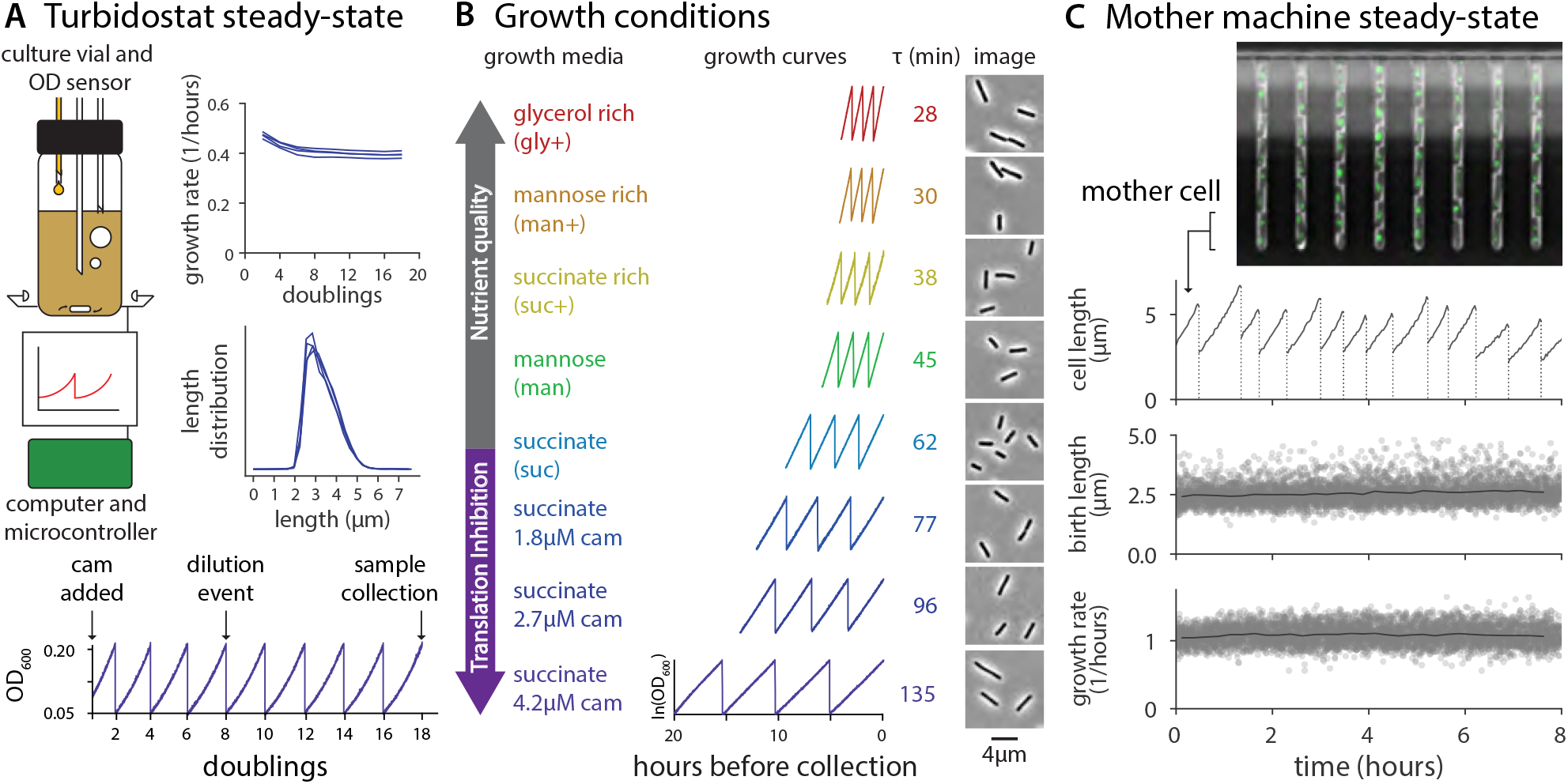
Population and single-cell methods to achieve steady-state growth. (A) Turbidostat experimental method and validation. Top left: In the multiplex turbidostat vial, the culture volume was maintained constant and cell concentration was monitored and adjusted automatically by infusing fresh medium. Aerobic conditions were ensured via bubbling and stirring. Top right: growth rate measurements were consistent between 5 and 20 doublings and cell length distributions were reproducible at sample collection. Data shown is from 4 repeats in succinate with 2.7 *µ*M chloramphenicol (cam). Bottom: A representative growth curve showing the timing of the addition of chloramphenicol, dilution events, and sample collection. Each dilution occurred when the culture reached OD_600_ 0.2 and was diluted to 0.05, allowing for two doublings during the growth interval. (B) Overview of population growth conditions and measurements. Growth media and their abbreviations for 5 different nutrient conditions, all of which are based on S7_50_. Glycerol rich, mannose, and succinate were selected for translation inhibition experiments (succinate shown here). Representative growth curves (final 8 doublings), average doubling time *τ*, and representative crops of images used for population sizing shown for each condition. (C) Single-cell experiments with the mother machine. Representative image showing cell-containing traps. Fluorescent signal is DnaN-mGFPmut2 (Figure **3**, Materials and methods). The growth in length (black lines) and division (dotted vertical lines) of a single mother cell is shown over 8 hours. Average birth length and growth rate (solid grey lines) of the single-cell measurements (grey scatter points) are in steady-state over the course of the experiment. Data shown is from mannose. Additional measurements for all conditions are presented in Extended Figure **1-1**.

To measure how long it takes for *B. subtilis* to reach physiological steady state, we measured growth rate continuously during time course experiments using our multiplex turbidostat. Growth rate generally stabilized after 6 generations, and the cell size distribution was reproducible (Figure **1**A). However, to be certain of steady-state growth, we typically waited for at least 14 doublings before sample collection in all our subsequent experiments. At collection, we split the culture for qPCR marker frequency and cell size measurement (see Table 5 for experimental conditions).

For single-cell measurements, we used the microfluidic mother machine to collect phase contrast and fluorescent timelapse images for at least 10 generations^7, 19^ (Figure **1**C). After analyzing all cell lives, we limited our data to the time interval in which all measured parameters equilibrated (Extended Figure **1-1**). A typical experiment produced data for around 2,500 cells (see Table 6 for experimental conditions).

### Growth law of cell size: *B. subtilis* size shows a positive but not exponential dependence on the nutrient-imposed growth rate

A foundational observation by Schaechter, Maaløe, and Kjeldgaard showed that the average cell size in *E. coli* increases exponentially with respect to the nutrient-imposed growth rate^2^. Previously, we investigated this ‘growth law of cell size’ in *E. coli* under various growth and cell cycle inhibition, and showed that the exponential relationship was a special case wherein the growth rate was the only experimental variable^3^. In *B. subtilis*, the Levin lab recently revisited the relationship between size and the nutrient-imposed growth rate, and found that the average cell size in *B. subtilis* increased with the growth rate at the population level^20^.

We extended our efforts in *E. coli* to *B. subtilis*. Using the multiplex turbidostat, we grew cells in 5 nutrient conditions with doubling times ranging between 28 and 62 minutes (Figure **1**B, Materials and methods; Table 5). Here, we use size interchangeably with volume, and consider volume to be proportional to dry mass^21^.

Figure **2**A shows the average cell size versus growth rate for the 5 different growth conditions. As expected, the average cell size increased with growth rate. However, the exponential dependence observed for *E. coli* was less clear in *B. subtilis*. This discrepancy in *B. subtilis* could be due to changes in the duration of replication (C period) and cell division (D period) in different nutrient conditions^3^.

**Figure 2:**
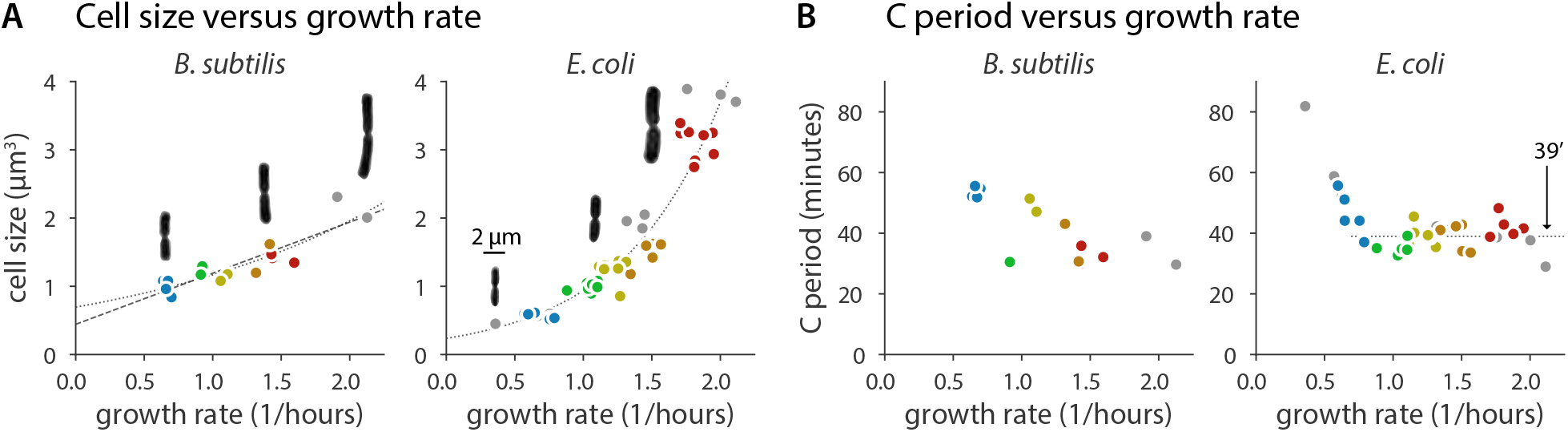
Population cell size and C period measurements in *B. subtilis* and *E. coli*. (A) Cell size increases with respect to growth rate in *B. subtilis* and *E. coli* under nutrient limitation. For *B. subtilis*, the relationship is not clearly exponential as it is for *E. coli* (dotted lines are linear regression fits of logarithm transformed data, dashed line is a linear regression fit). Representative images of cells during division show change in aspect ratio as a function of growth rate. Length and width measurements presented in Extended Figure **2-1**. (B) C period measurements with respect to growth rate in *B. subtilis* and *E. coli* under nutrient limitation. For *E. coli*, C is approximately constant at 39 minutes (horizontal dotted line) for doubling times faster than 60 minutes (*λ* = 0.69). That constancy is less clear for *B. subtilis*, though single-cell data shows that C+D is proportional to generation time (Extended Figure **4-1**). *B. subtilis* growth media are colored as in Figure **1**B with additional LB data in grey. *E. coli* data is previously published work^3^; Red is synthetic rich, orange is glucose with 12 amino acids, yellow is glucose with 6 amino acids, green is glucose, and blue is glycerol, with additional conditions in grey.

We thus measured the population average C period of *B. subtilis* employing qPCR marker frequency analysis^3, 10, 22^. Both species exhibited a similar maximum replication speed (approximately 40 minutes for C period), but our data indeed do not indicate C period is strictly constant in fast growth (Figure **2**B).

Unfortunately, despite extensive efforts, we were unable to reliably measure the D period in *B. subtilis* from the population samples as we had done previously for *E. coli*^3^. The main issue was consistency of fluorescence labeling of the DNA required for flow or image cytometry. Our results were variable from experiment to experiment, and protocol to protocol. We therefore concluded that the measurement of D period using population methods is not as reliable as needed to test the growth law of cell size in *B. subtilis*, a cautionary reminder in interpreting previous measurements in *B. subtilis*. For these reasons, we set out to measure the *B. subtilis* cell cycle explicitly at the single-cell level.

### Single-cell determination of cell cycle parameters in *B. subtilis*

We employed a functional replisome protein fused with a fluorescent marker, DnaN-mGFPmut2, to measure cell cycle progression in single cells^7, 23^ (Materials and methods). In *B. subtilis*, the replisomes from the two replication forks of a replicating chromosome are often colocalized, thus most foci represent a pair of replisomes^24^.

Figure **3**A and B show representative cells from two growth conditions, succinate and glycerol rich, respectively. In the slower growth condition (succinate), cells were normally born with one replicating chromosome. Replication initiation begins synchronously in the mother cell for two chromosomes. At that time, the origins are located towards the cell poles. Replication proceeds through cell division, at which point the replication forks reside near the midcell of the newly born cell. Chromosome segregation is concurrent with replication. By the time the replication forks reach the terminus region, which is still at the midcell, the previously duplicated origins have already migrated to the cell poles^25^.

**Figure 3:**
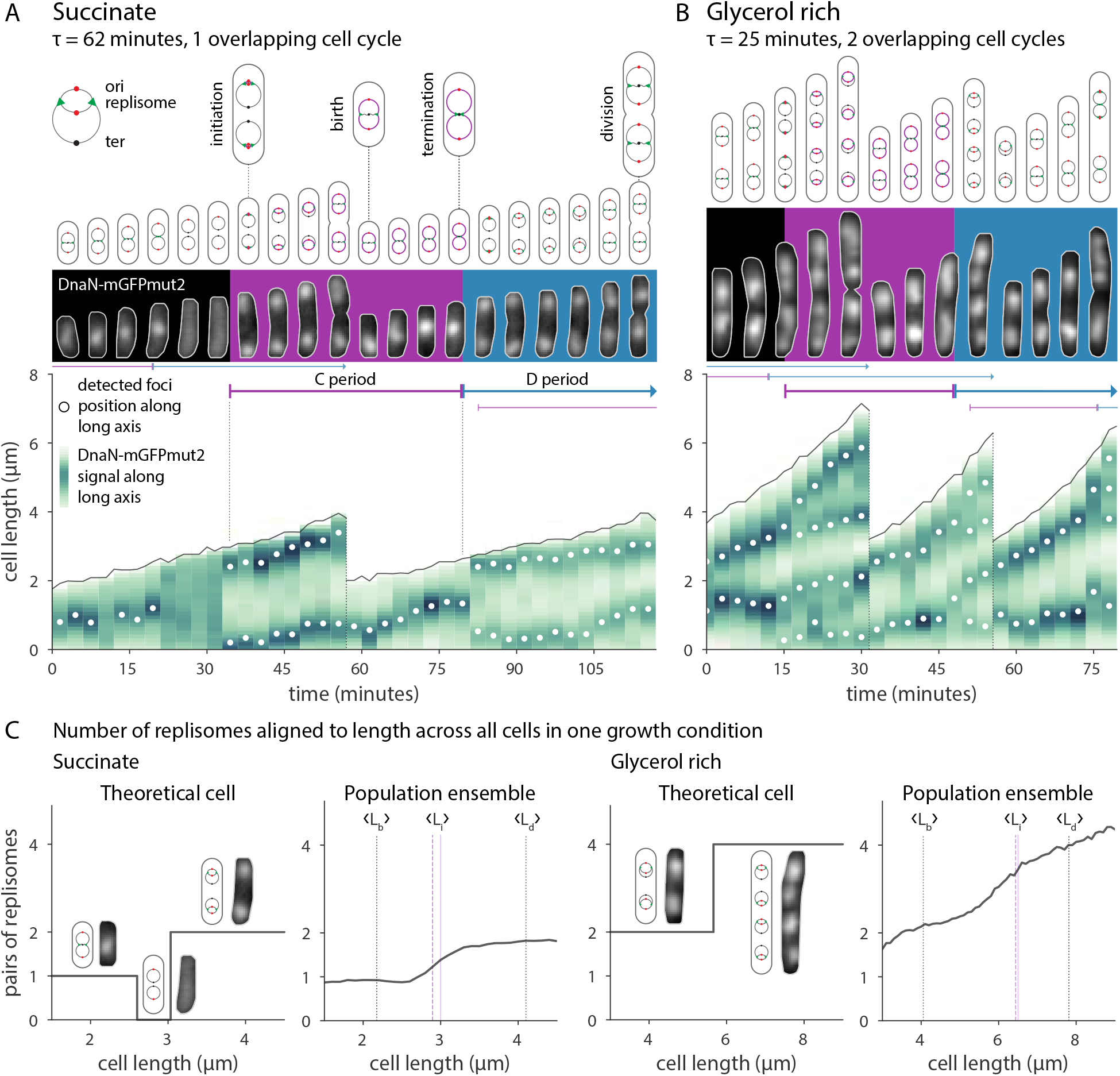
Single-cell growth and cell cycle progression in *B. subtilis*. (A) Typical cell cycle progression of *B. subtilis* in slower growth media. All panels show the same two cells. Top: Chromosome configuration and key cell events. Middle: Fluorescent images of DnaN-mGFPmut signal. Grey outlines are from the segmented phase contrast image. Purple and blue backgrounds indicate C and D periods corresponding to the second division. Bottom: Processed image data represented as a cell lineage trace. Cell length and division are indicated by the solid and dotted black lines, respectively. Vertical green bars are the DnaN-mGFPmut2 signal summed along the long axis of the cell, with white circles showing foci position. The single-cell initiation size, C period, and D period, are determined manually from these traces. (B) Typical cell cycle progression of *B. subtilis* in faster growth media. (C) Ensemble method to determine cell cycle parameters. Left: In succinate, a theoretical cell is born with one pair of replisomes. It may briefly contain no active replisomes upon termination, and then contain two pairs of replisomes as the two complete chromosomes begin replication. The length at which the number of replisome pairs increases corresponds to the initiation size. Across all cells, the average number of replisome pairs transitions from one to two at the population’s initiation size. The average initiation length <L_i_> as determined from the cell traces (dashed purple line) agrees with the ensemble estimate (solid purple line). The average birth <L_b_> and division length <L_d_> of the population are shown as dotted vertical lines. Right: In glycerol rich, cells transition from two to four pairs of replisomes. Ensembles for all conditions are available in Extended Figure **3-1**.

While overlapping cell cycles are common even at slower growth, cells rarely exhibit multifork replication. Multifork replication indicates initiation begins before the previous termination event completes. Instead, *B. subtilis* normally initiates when the cell contains complete, homologous chromosomes where the copy number is a power of two. In fact, replication initiation often proceeds immediately after the previous termination event. This may be due to the role of YabA in *B. subtilis* replication initiation control, which ties DnaA activity to DnaN availability^26, 27^. Comparatively, multifork replication is common in *E. coli*, where Hda is thought to play a similar but mechanistically distinct role in reducing initiation potential during ongoing replication^7, 28^.

In faster growth conditions (glycerol rich), cells are often born with two replicating chromosomes. However, the relative variability between division size and C+D was greater in this rich condition. This means that a substantial fraction of the population were still born with one replicating chromosome (Extended Figure **6-2**). Moreover, transient filamentation and asymmetrical septation are more common in fast growth conditions, leading to cells born with a number of replicating chromosomes which are not a power of two.

### Complementary, ensemble determination of cell cycle parameters in *B. subtilis*

The main advantage of the single-cell approach is that it allows for direct comparison of the relationships between growth parameters, providing mechanistic insights^6^. However, it can be difficult to determine the cell cycle parameters manually, particularly when the foci are clumped or the signal is weak. This is especially true in faster growth conditions. To ensure an unbiased analysis of the cell cycle, we also employed an “ensemble method” to extract cell cycle parameters^11^ (Figure **3**C). We used the foci count at a given size as a proxy for the replication state (Materials and methods). This method produces data similar to the original schematics used by Helmstetter and Cooper when first elucidating the *E. coli* cell cycle^29^.

For all but the slowest growth conditions, the measured average number of foci monotonically increases because initiation almost immediately follows termination as discussed above. Unlike a theoretical single cell, the ensemble plots do not display a strict step-like behavior; we interpret this as variability in the initiation size. Ensemble plots for all conditions, along with the foci localization patterns, are presented in Extended Figure **3-1**. This measurement is in good agreement with the average initiation size as measured from individual cells. We used these complementary methods to test whether the initiation size is invariant in *B. subtilis* as in *E. coli*^3^.

### Invariance of initiation size: *B. subtilis* initiates at a fixed cell size per *ori*

In *E. coli* and *S. typhimurium*, the concept of a conserved initiation size was first explained by Donachie as a consequence of the growth law of cell size and the constant C+D^2, 8, 29^. The upshot is that, at a fixed size per origin (*ori*), all origins of replication fire simultaneously. Recent highthroughput works at both single-cell and population levels have conclusively shown that the early insight by Donachie was correct^3, 7, 11^. In fact, the initiation size per *ori* is invariant not only across nutrient conditions, but also under antibiotic inhibition and genetic perturbations^3^.

The constancy of initiation size in *B. subtilis* has previously been tested by several groups at the population level under nutrient limitation conditions^9, 10, 30^. We further measured the initiation size using single-cell methods under nutrient limitation and translational inhibition. We found that the initiation size per *ori* in *B. subtilis* is indeed invariant across conditions, even for individual cells (Figure **4**A).

This constant initiation size is in stark contrast to the varying C period under different growth conditions (Extended Figure **4-1**A). In fact, initiation size is one of the least variable physiological parameters along with septum position and width (Extended Figure **6-2**). The single-cell approach also allowed us to measure the correlations between all growth and cell cycle parameters. The initiation size is only weakly correlated with other measured parameters (Extended Figure **6-3**).

**Figure 4:**
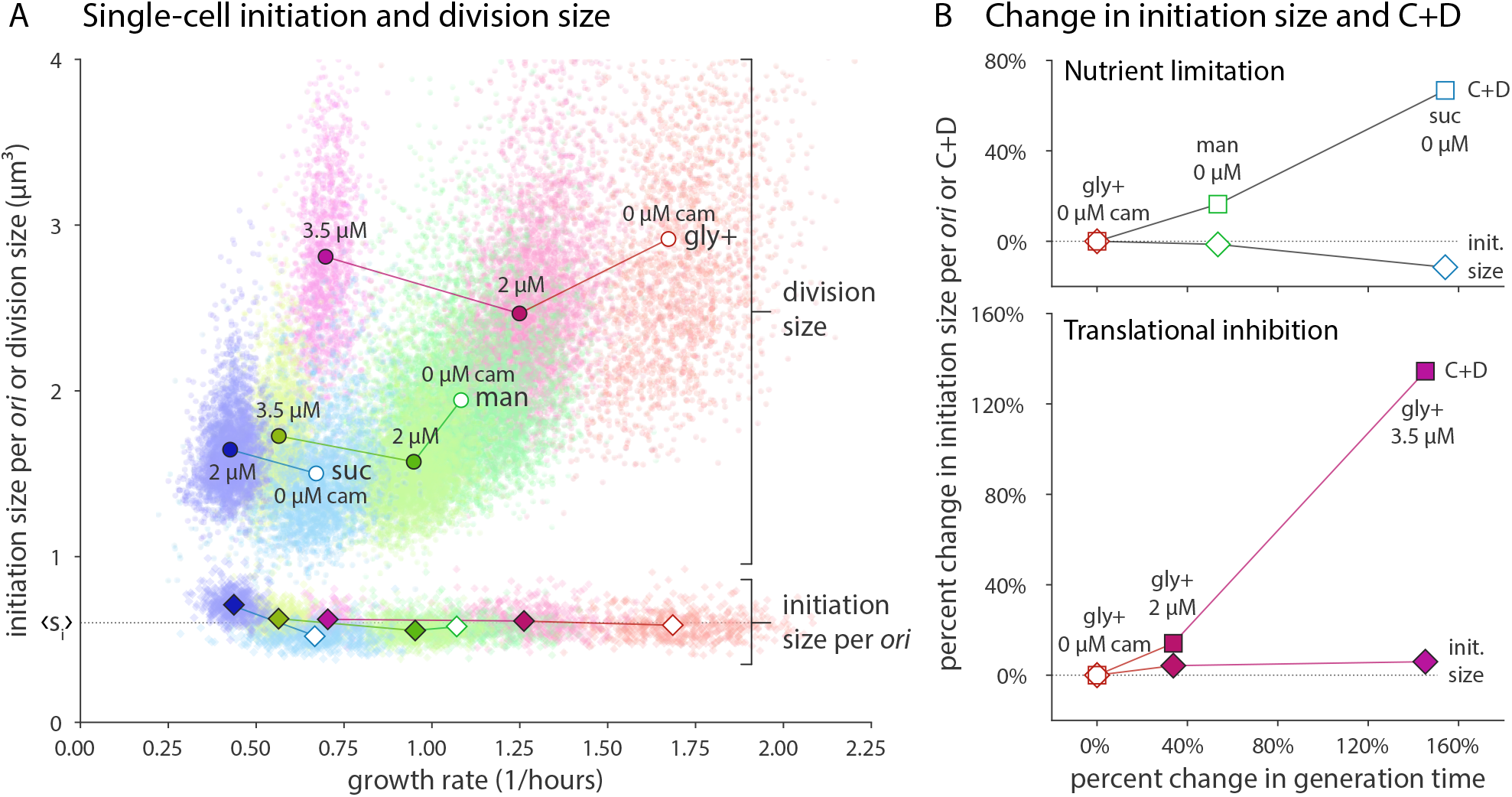
Initiation size is invariant in *B. subtilis* during steady-state growth. (A) Single-cell initiation size per *ori* s_i_ is condition independent. Division size (circles) changes dramatically under both nutrient limitation and translational inhibition. The corresponding initiation size per *ori* (diamonds) collapses onto a constant value, <s_i_>, across all conditions (dotted horizontal black line). This holds for both single cells (scatter points) and population averages (solid symbols). Growth media are colored as in Figure **1**B with the amount of chloramphenicol indicated (0 *µ*M chloramphenicol with empty symbols, lines connect the same growth media with and without chloramphenicol). (B) C+D is condition dependent and increases with generation time. Top: Increase in generation time under nutrient limitation causes an increase in the population average C+D (squares) while initiation size minimally changes. Bottom: A similar pattern is seen under translational inhibition. For both plots, measured parameters are compared to the fastest growth condition. Single-cell C+D data presented in Extended Figure **4-1** and Extended Figure **6-2**.

These observations are consistent with a threshold model for replication initiation^3, 7, 31^. Within that framework, initiator molecules accumulate proportional to the growth rate. This mechanism is enacted in single cells and is in turn apparent at the population level.

### Initiation size is invariant even during nutrient shifts at the single-cell level

Because the constant initiation size was implemented by individual cells in the previous steady-state experiments, we wondered how cells would behave in a changing environment. Nutrient shift experiments have provided important insight into the coordination of biosynthesis and the cell cycle^32–34^. We revisited this paradigm at the single-cell level, shifting cells from minimal media (*τ* = 65 min) to rich conditions (*τ* = 30 min) and back again (Extended Figure **5-1**). By using the mother machine, we could add and remove nutrients immediately while measuring the cell cycle and all other physiological parameters (Materials and methods).

The most drastic results occurred upon shift-down (Figure **5**). When nutrient supplements were removed, growth immediately paused. The crash in growth rate caused a drastic increase in generation and cell cycle time for cells which experienced the shift-down. Replicating chromosomes were stalled and division ceased. Strikingly, the growth pause led to an absence of initiation events until after cells restarted elongation and attained the requisite initiation size. Thus individual cells maintained a constant initiation size through the transition. Division also resumed after growth recommenced, but at a smaller size commensurate with the postshift-down growth rate. A constant C+D period is not maintained during this time (Extended Figure **5-1**).

**Figure 5:**
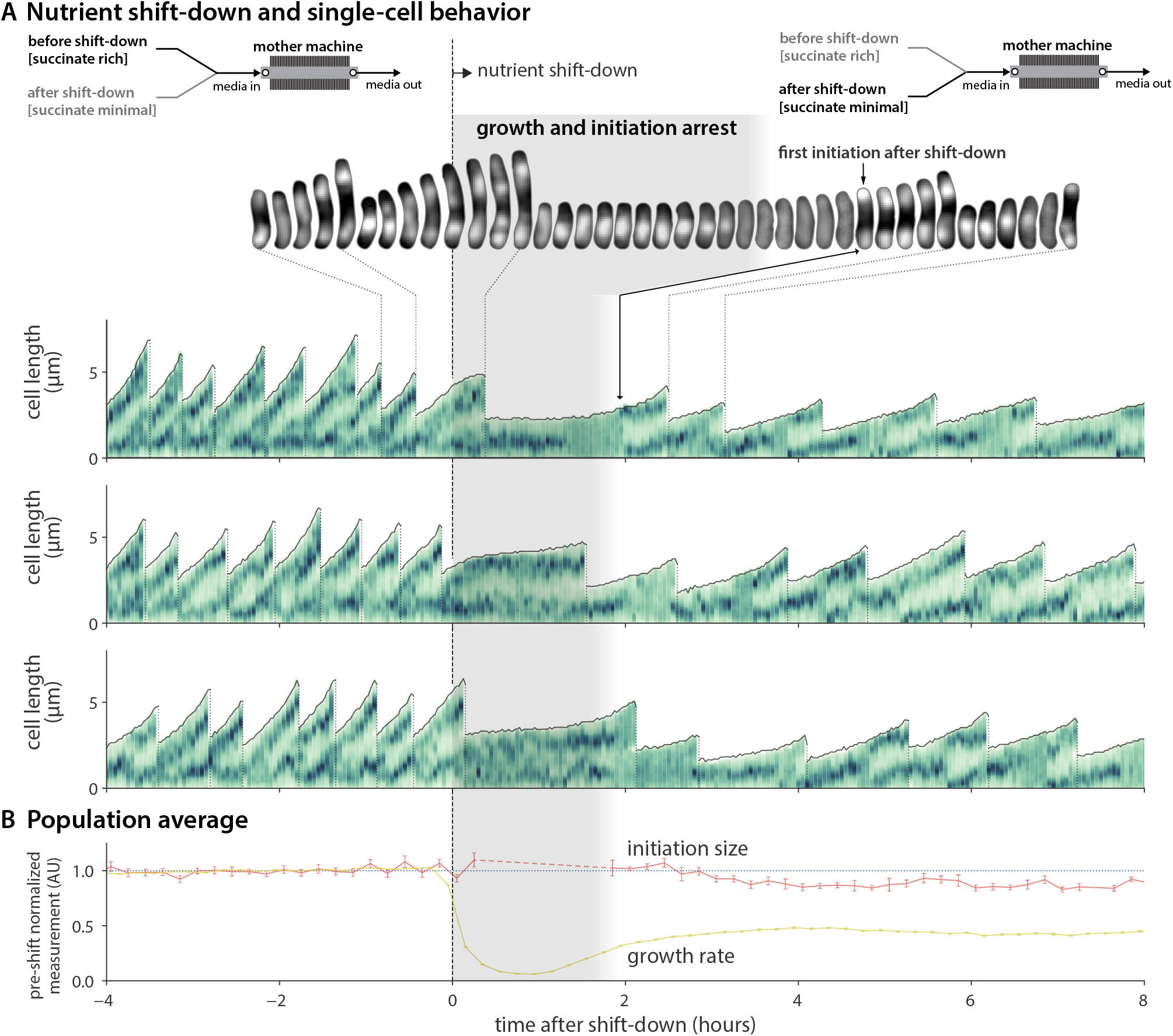
Initiation size is invariant in *B. subtilis* during shift-down. (A) Behavior of single cells undergoing shift-down from succinate rich to succinate minimal at time zero. Upon shift-down, cells pause growth and initiation for 1-2 hours. Media shift is achieved via a Y-valve upstream of the mother machine inlet. Top: Fluorescent images of cell lineage during shift-down. Bottom: Representative traces of 3 lineages. Representation is as in Figure **3**, with the density of the vertical green bars representing the intensity of the DnaN-mGFPmut2 signal along the long-axis of the cell. (B) Population average behavior during shift-down. Growth rate is the instantaneous elongation rate (6 minute time step). Initiation size per *ori* is plotted against the initiation time. Each measurement is normalized by their respective mean in the 4 hours before shift-down. Lines connect the 12 minute binned mean, and error bars are the standard error of the mean. Minimum bin size is 5. Dashed line in average initiation size signifies gap in initiation events after shift-down. n=3,160 cells (752 with initiation size). Entire time course showing shift-up and shift-down is available in Extended Figure **5-1**.

The decoupling of initiation and division supports the idea that they are controlled by independent threshold mechanisms^7^. That is, the cell builds up a pool of dedicated molecules for each task to a certain level^7, 12, 35–37^. For initiation, this threshold and the accumulation rate is conserved across growth conditions. For division, the threshold or the accumulation rate is set by the growth condition^38^. In the generation after shift down, cells grow much more slowly and therefore accumulate threshold molecules at a similarly depressed rate. As a result, both initiation and division are delayed. For division, active degradation or antagonization of FtsZ could further hinder the triggering of constriction^39, 40^.

### *E. coli* and *B. subtilis* change cell shape differently under different growth conditions but maintain a constant initiation size

One of the major differences between *E. coli* and *B. subtilis* is their shape under different nutrient conditions. Data from our lab and others have shown that the aspect ratio of *E. coli* is nearly constant (approximately 4) under different nutrientimposed growth rates^3, 41^. By contrast, the average width of *B. subtilis* remains relatively constant (Extended Figure **2-1**)^9, 42^.

Nevertheless, for initiation control in *B. subtilis*, we find that volume per *ori* is more conserved than the length per *ori* at initiation. While we find length to be a good proxy for initiation size under nutrient limitation, our data show that chloramphenicol treatment decreases cell width in *B. subtilis*. Thus, when comparing across all growth conditions, only the initiation volume is constant (Extended Figure **4-1**B-D).

### *B. subtilis* is both a division adder and an initiation adder

As previously reported, *B. subtilis* achieves size homeostasis by following the adder principle^6^. We recently showed that *B. subtilis*, along with *E. coli*, are also initiation adders; the size added per *ori* between successive initiation events is constant with respect to initiation size^7^. We further tested those results here under additional growth conditions and translational inhibition (Extended Figure **4-2**). We find that, for division, our data is best described by the adder principle. However, we note that when going from faster to slower growth condition, the slope becomes slightly negative. This may be due to active degradation or inhibition of FtsZ assembly or other key division proteins^7, 39, 40^. For initiation, we again find that our data is best described by the adder principle. Importantly, the added size between initiation and added size between division are uncorrelated (Extended Figure **6-3**), consistent with initiation and division being controlled by separate threshold mechanisms^7^. While both processes are tied to global biosynthesis, this indicates minimal crosstalk between the two in steady-state conditions.

### *B. subtilis* and *E. coli* share the same hierarchy of physiological parameters

The coefficient of variation (CV) of a distribution of a physiological parameter is often interpreted as the tightness of the underlying biological control^43^. We extended previous analysis to include the cell cycle related parameters C period, D period, initiation size, and added initiation size for both *B. subtilis* and *E. coli*. We found that both evolutionarily distant organisms share the same order of their physiological parameters in terms of CV (Figure **6**). Width, septum position, initiation size, and growth rate are the tightest of the parameters. D period is significantly more variable than C period, and they are inversely correlated. In fact, the CV of a particular physiological parameter is extremely similar across growth conditions, species, and strains (Extended Figure **6-1**).

**Figure 6:**
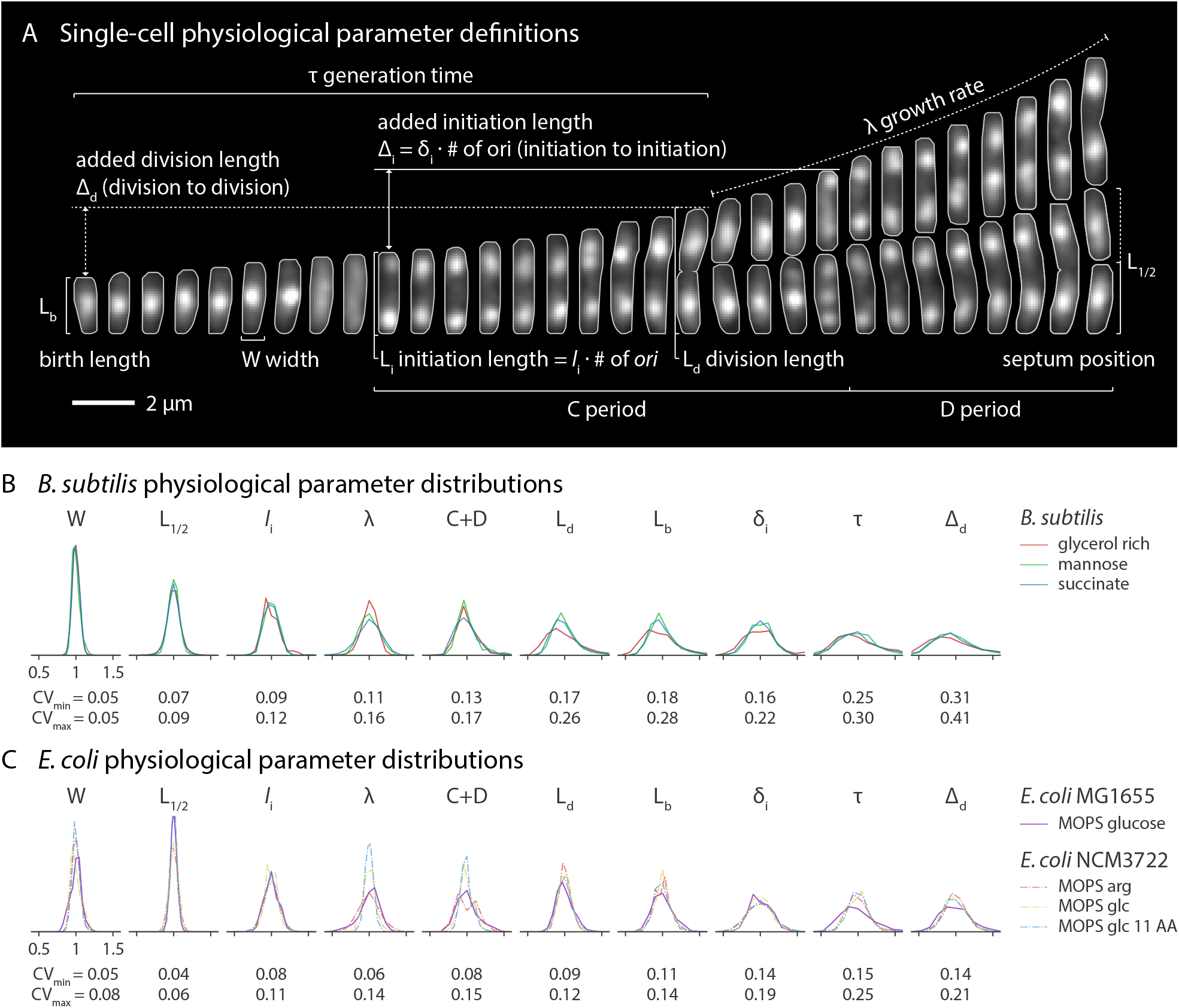
*B. subtilis* and *E. coli* share the same hierarchy of physiological parameters. (A) Single-cell physiological parameter definitions as determined from time-lapse images. Cells are *B. subtilis* growing in mannose (*τ* = 38 minutes). Fluorescent signal is DnaN-mGFPmut2 and grey outlines are from the segmented phase contrast image. Picture interval is 3 minutes. (B) *B. subtilis* parameter distributions are shown in order of ascending coefficient of variation (CV). Parameters are normalized by their mean. The range of CVs for each parameter is shown below the distributions. Note that length based parameters are shown here; their volume equivalents have slightly higher CVs due to variability in width. (C) Distribution of the same measurements in *E. coli* display the same CV hierarchy. Data from *E. coli* NCM3722 grown in MOPS arginine (arg), glucose (glc) and glucose + 11 amino acids (glc 11 AA) are from previously published work^7^.

Ultimately, the CV of the physiological parameters is the manifestation of molecular regulatory mechanisms. Classically, *B. subtilis* and *E. coli* provide excellent examples of both homologous and non-homologous versions of such mechanisms. For example, major protein players controlling replication and division, such as DnaA and FtsZ, are conserved in these and most other prokaryotes^44, 45^. However, the regulation of those molecules in *B. subtilis* and *E. coli* is unique^46–48^. More generally, the two species often use unrelated mechanisms to achieve the same regulatory goal^48, 49^. Because of their phylogenetic distance, the uncanny agreement between the CVs of their physiological parameters suggests an evolutionary ancient control framework shared by these organisms.

## Summary and Outlook

We have shown that *B. subtilis* and *E. coli*, despite their historical separation across the Gram stain divide, share extremely similar fundamental physiological behavior (Figure **7**). Under a wide range of nutrient and growth inhibition conditions, both species base their chromosome replication in a constant initiation size. Impressively, this constant initiation size is imposed even during dynamic growth transitions. This is consistent with a threshold mechanism and constant production of cell cycle initiator proteins for initiation and division timing control, thus maintaining size homeostasis with the adder principle^7^.

**Figure 7:**
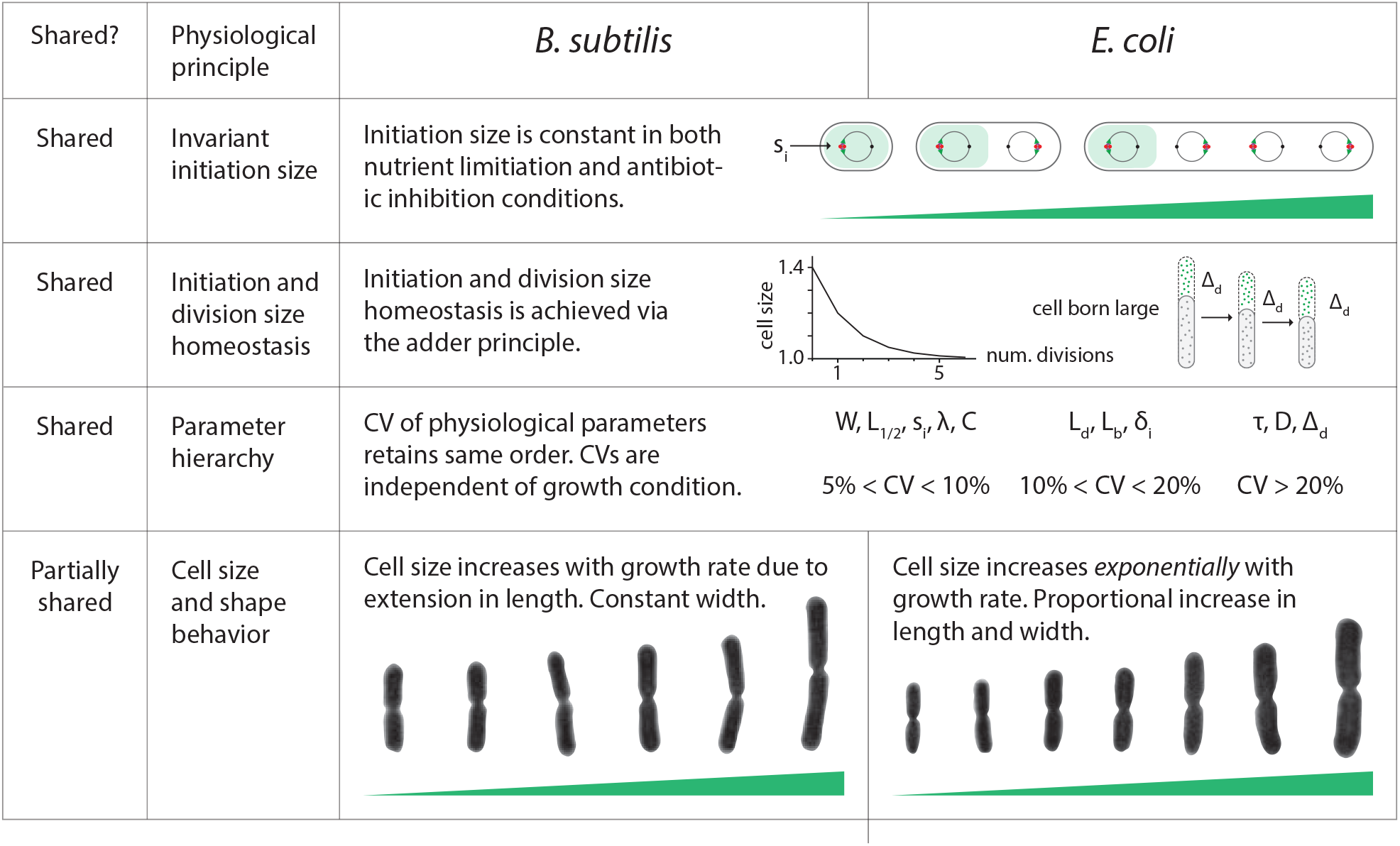
*B. subtilis* and *E. coli* comparative summary.

As with *E. coli*, DnaA and FtsZ are among the key proteins responsible for the initiation and division threshold mechanisms in *B. subtilis*, respectively^7, 13, 50^. The view that global biosynthesis fundamentally controls their production, and thus the replication and division rate, is still compatible with the idea that additional levels of regulation modulate or coordinate their activity in certain situations^46, 51^. It is unclear whether these additional mechanisms have evolved to increase replication and division fidelity during steady-state or are more important in dynamic environments. More single-cell shift experiments with mutant or even minimal genome cells will help reveal the importance of redundant regulatory systems.

These deep similarities between *B. subtilis* and *E. coli* speak to a conserved control framework which both species use to coordinate growth, DNA replication, and division. In doing so, they ensure life’s essential demand of physiological homeostasis. In the end, it is unclear if this framework is the result of parallel or convergent evolution. In order to better address this question, more quality single-cell data is needed from diverse prokaryotes. In either case, the existence of a shared control framework underscores its efficacy, providing an intriguing avenue for the development of synthetic organisms.

## Acknowledgements

This work was supported by the Paul G. Allen Family Foundation, Pew Charitable Trust, NSF CAREER grant MCB-1253843, and NIH grant R01 GM118565-01 (to S.J.).

## Author Information

The authors declare no competing financial interests. Correspondence and requests for materials should be addressed to S.J.

## Materials and methods

### Strains

We used *B. subtilis* strains in the 3610 background with mutations to confer non-motility and reduce biofilm formation^6^. The background strain contained *comI*(Q12L) to confer competence^52^. We used an inducible *lytF* construct to prevent chaining^18^. For mother machine experiments in which replisomes were tracked, we used *dnaN-mGFPmut2*^23, 24^. Strain construction was performed using single crossover plasmid recombination or double crossover recombination from genomic DNA^53^.

For *E. coli*, we used a K-12 MG1655 strain containing a functional *dnaN-YPet* construct^54^. Strain genotypes for both species are provided in Table 2.

**Table 1:** Strain information.

| *B. subtilis* strains | Genotype | Notes |
| --- | --- | --- |
| BS15 | 3610 comI(Q12L) hag::MLS(R)::MLS(S)<br>amyE::[Phyperspank-lytF spcR] epsH::tet | Gift from Petra Levin; Used for<br>turbidostat experiments. |
| BS45 | 3610 comI(Q12L) motAB::Tn917<br>amyE::[Physpank-lytF kan] epsH::tet<br>dnaN::[dnaN-gfp spec] | This study; Used for microfluidic<br>and turbidostat experiments. |
| *E. coli* strains |  |  |
| SJ1724 | K-12 MG1655 dnaN::[dnaN-yPet kan]<br>hupA::[hupA-mRuby2 FRT-cat-FRT] | This study; Used for microfluidic<br>experiments. |

**Table 2:**
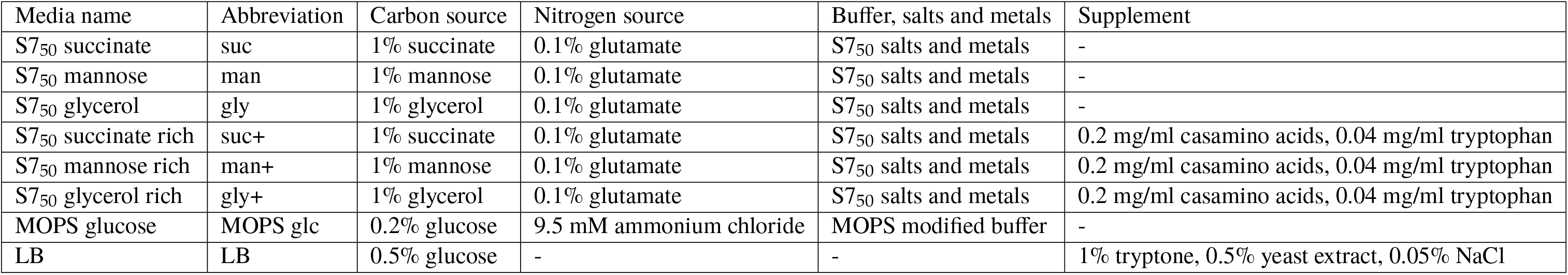
Growth media.

| Media name | Abbreviation | Carbon source | Nitrogen source | Buffer, salts and metals | Supplement |
| --- | --- | --- | --- | --- | --- |
| S7<sub>50</sub> succinate | suc | 1% succinate | 0.1% glutamate | S7<sub>50</sub> salts and metals | - |
| S7<sub>50</sub> mannose | man | 1% mannose | 0.1% glutamate | S7<sub>50</sub> salts and metals | - |
| S7<sub>50</sub> glycerol | gly | 1% glycerol | 0.1% glutamate | S7<sub>50</sub> salts and metals | - |
| S7<sub>50</sub> succinate rich | suc+ | 1% succinate | 0.1% glutamate | S7<sub>50</sub> salts and metals | 0.2 mg/ml casamino acids, 0.04 mg/ml tryptophan |
| S7<sub>50</sub> mannose rich | man+ | 1% mannose | 0.1% glutamate | S7<sub>50</sub> salts and metals | 0.2 mg/ml casamino acids, 0.04 mg/ml tryptophan |
| S7<sub>50</sub> glycerol rich | gly+ | 1% glycerol | 0.1% glutamate | S7<sub>50</sub> salts and metals | 0.2 mg/ml casamino acids, 0.04 mg/ml tryptophan |
| MOPS glucose | MOPS glc | 0.2% glucose | 9.5 mM ammonium chloride | MOPS modified buffer | - |
| LB | LB | 0.5% glucose | - | - | 1% tryptone, 0.5% yeast extract, 0.05% NaCl |

### Growth media and experimental conditions

For *B. subtilis*, we used S7_50_ medium with different carbon sources and supplements. Importantly, we included additional iron(III) chloride and trisodium citrate. The latter acts as a siderophore for *B. subtilis*, and without it our strain cannot grow in the mother machine^55^. To make rich conditions, we added 2 mg/mL casamino acids and 0.04 mg/mL tryptophan. For *E. coli*, we used MOPS glucose medium. Turbidostat and mother machine experiments used the same media with the following addition: bovine serum albumin was added at 0.5 mg/mL during mother machine experiments in order to reduce cell adherence to surfaces inside the device. Tables 3 and 4 provide detailed information on media composition.

**Table 3:** Media components.

| Component | Concentration |
| --- | --- |
| S7<sub>50</sub> salts and metals |  |
| MOPS | 50 mM |
| ammonium sulfate | 1 mM |
| potassium phosphate monobasic | 5 mM |
| magnesium chloride | 2 mM |
| calcium chloride | 0.7 mM |
| manganese(II) chloride | 50 μM |
| zinc chloride | 1 μM |
| iron(III) chloride | 55 μM |
| thiamine hydrochloride | 1 mM |
| hydrogen chloride | 20 μM |
| trisodium citrate | 50 μM |
| MOPS modified buffer |  |
| MOPS | 40 mM |
| tricine | 4 mM |
| iron(III) sulfate | 0.1 mM |
| sodium sulfate | 0.276 mM |
| calcium chloride | 0.5 μM |
| magnesium chloride | 0.525 mM |
| sodium chloride | 50 mM |
| ammonium molybdate | 3 nM |
| boric acid | 0.4 μM |
| cobalt chloride | 30 nM |
| cupric sulfate | 10 nM |
| manganese(II) chloride | 80 nM |
| zinc sulfate | 10 nM |
| potassium phosphate monobasic | 1.32 mM |

**Table 4:**
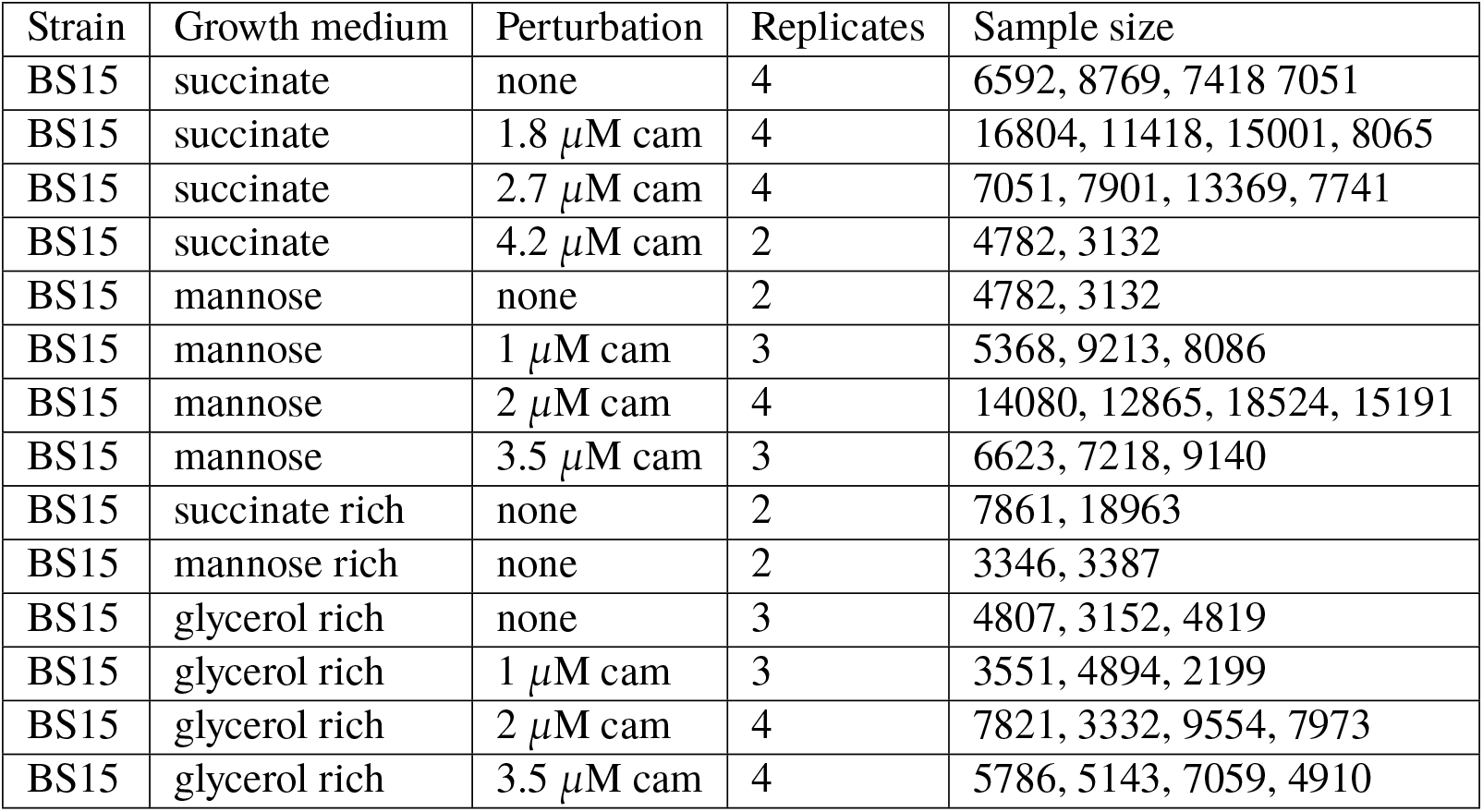
Turbidostat experimental conditions.

| Strain | Growth medium | Perturbation | Replicates | Sample size |
| --- | --- | --- | --- | --- |
| BS15 | succinate | none | 4 | 6592, 8769, 7418 7051 |
| BS15 | succinate | 1.8 μM cam | 4 | 16804, 11418, 15001, 8065 |
| BS15 | succinate | 2.7 μM cam | 4 | 7051, 7901, 13369, 7741 |
| BS15 | succinate | 4.2 μM cam | 2 | 4782, 3132 |
| BS15 | mannose | none | 2 | 4782, 3132 |
| BS15 | mannose | 1 μM cam | 3 | 5368, 9213, 8086 |
| BS15 | mannose | 2 μM cam | 4 | 14080, 12865, 18524, 15191 |
| BS15 | mannose | 3.5 μM cam | 3 | 6623, 7218, 9140 |
| BS15 | succinate rich | none | 2 | 7861, 18963 |
| BS15 | mannose rich | none | 2 | 3346, 3387 |
| BS15 | glycerol rich | none | 3 | 4807, 3152, 4819 |
| BS15 | glycerol rich | 1 μM cam | 3 | 3551, 4894, 2199 |
| BS15 | glycerol rich | 2 μM cam | 4 | 7821, 3332, 9554, 7973 |
| BS15 | glycerol rich | 3.5 μM cam | 4 | 5786, 5143, 7059, 4910 |

For both turbidostat and mother machine experiments, chloramphenicol was added at concentrations between 1- and 4.2 *µ*M during translational inhibition experiments. All experiments were performed at 37^◦^C in a climate controlled environmental room which housed the multiplex turbidostat and all optical components (Darwin Chambers Company, MO). Tables 5 and 6 enumerate experimental conditions and sample size for turbidostat and mother machine experiments, respectively.

**Table 5:**
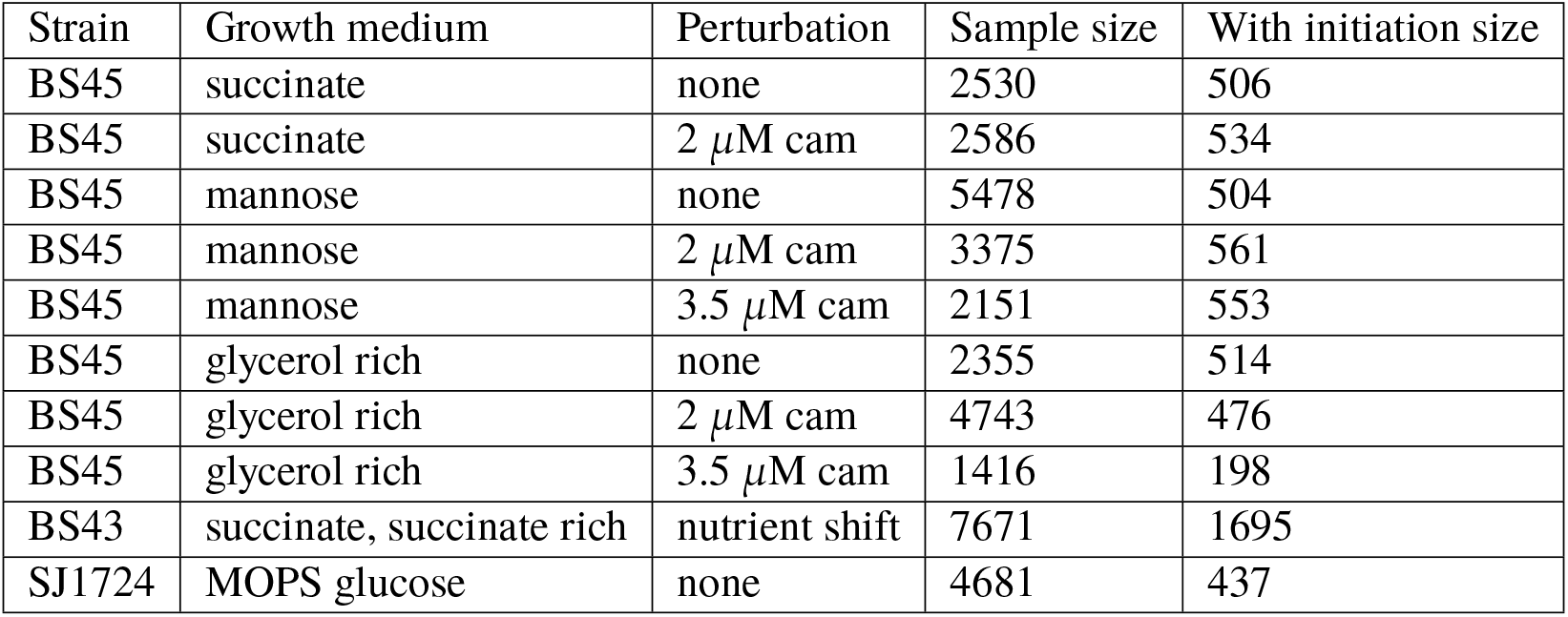
Mother machine experimental conditions.

| Strain | Growth medium | Perturbation | Sample size | With initiation size |
| --- | --- | --- | --- | --- |
| BS45 | succinate | none | 2530 | 506 |
| BS45 | succinate | 2 μM cam | 2586 | 534 |
| BS45 | mannose | none | 5478 | 504 |
| BS45 | mannose | 2 μM cam | 3375 | 561 |
| BS45 | mannose | 3.5 μM cam | 2151 | 553 |
| BS45 | glycerol rich | none | 2355 | 514 |
| BS45 | glycerol rich | 2 μM cam | 4743 | 476 |
| BS45 | glycerol rich | 3.5 μM cam | 1416 | 198 |
| BS43 | succinate, succinate rich | nutrient shift | 7671 | 1695 |
| SJ1724 | MOPS glucose | none | 4681 | 437 |

**Table 6:**
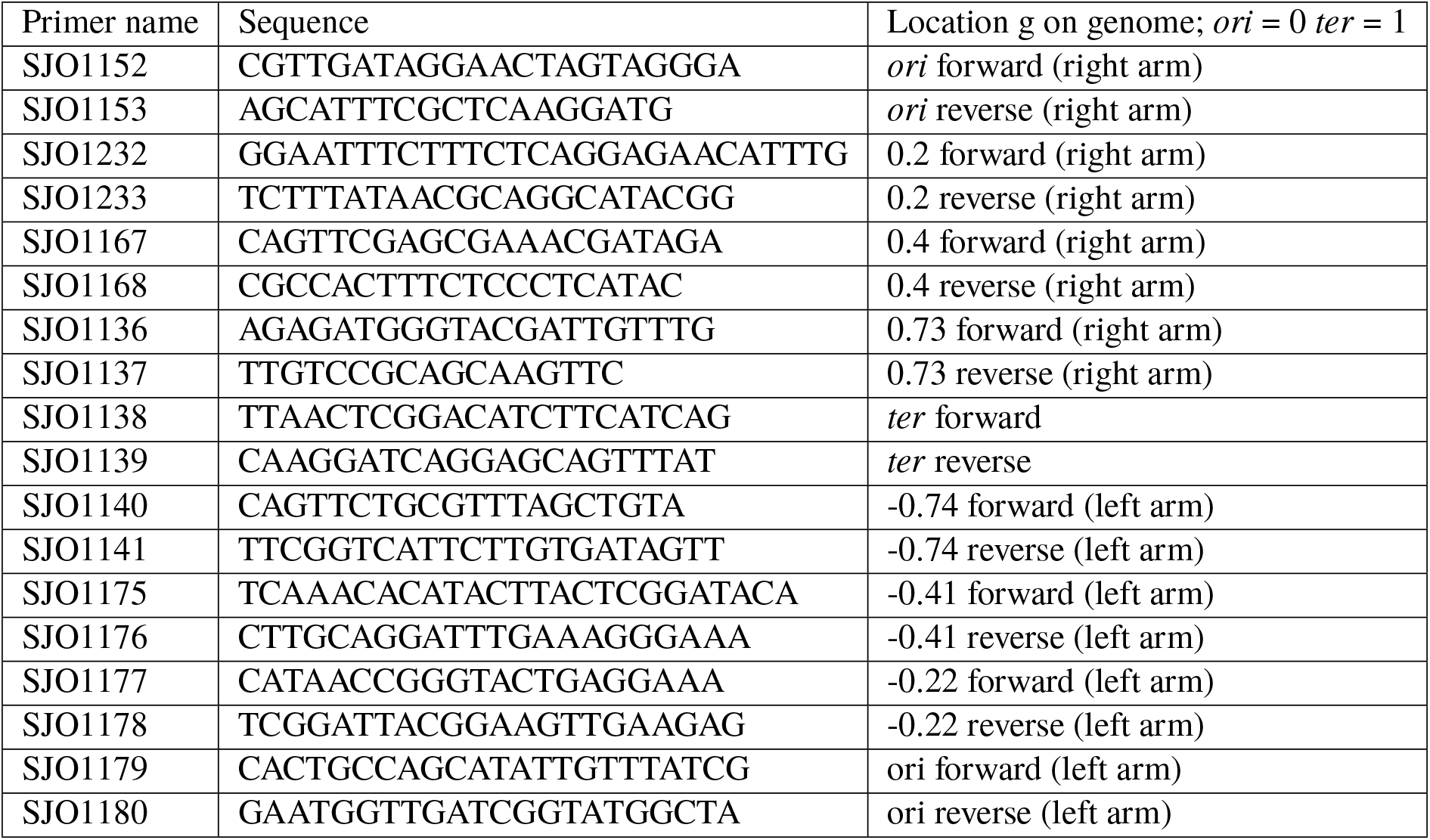
qPCR primers.

### Microscopy configuration

We performed phase contrast and fluorescent imaging on a Nikon Ti-E inverted microscope with Perfect Focus (PFS) and an LED transmission light source, controlled by Nikon Elements. For turbidostat experiments we used a PFS 2, CoolLED pE-100, 60X 1.4 NA Ph3 oil immersion objective (Nikon CFI Plan Apo DM Lambda 60X Oil), and Andor Technology Neo sCMOS camera. For fixed cell phase contrast imaging, we used exposure times between 50-100 ms and 100% transmission power.

For mother machine experiments, we used a PFS 3, Sutter Instruments TLED, 100X 1.45 NA Ph3 oil immersion objective (Nikon CFI Plan Apo DM Lambda 100X Oil), Photometrics Prime 95B sCMOS camera, and Coherent Obis laser 488LX for epifluorescent illumination. For laser epifluorescent illumination, we inserted a rotating diffuser in the optical train to reduce speckle. We also reduced the camera sensor region of interest to flatten the fluorescent illumination profile. We used a Chroma filter cube with dichroic mirror ZT488rdc and emission filter ET252/50m. For live cell phase contrast imaging, we used a 30 ms exposure time at 100% transmission power at an interval of 1.5 minutes. For fluorescent imaging, we used a 25 or 50 ms exposure time at 25% power at an interval of 3 minutes. This weak illumination minimized physiological effects due to phototoxicity on the cell and allowed for steady-state behavior over many hours.

### Turbidostat cell preparation and sample collection

We grew all pre-cultures at 32^◦^C or 37^◦^C in a water bath shaker at 260 rpm. Seed cultures were inoculated into 1-3 mL LB medium from a single colony from an agar plate, streaked no more than 2 days before use. Cells were grown for several hours then diluted 1,000-fold into the target media without antibiotics and grown until OD_600_ 0.1. If multiple back dilution rounds were needed to control experimental timing, they were done such that cells did not enter stationary phase. The culture was then inoculated into each turbidostat vial with or without antibiotics to the target OD_600_ 0.05. Cultures grew for a minimum of 14 doublings to ensure steady-state conditions upon sample collection. For some conditions, cells adhered to the glass culture vial, evidence of residual biofilm activity we observed as changes in growth rate over the time course. In these cases, the sample was transferred to a clean glass vial at the end of the experiment for at least 1 additional doubling from which the growth rate was determined.

We collected samples for cell size and cell cycle measurements at OD_600_ 0.2. Approximately 20 mL of cell culture was immediately put on ice to arrest growth. The culture was then split and pelleted, frozen, or fixed according to the subsequent measurement protocol. Our turbidostat design and function has been previously described^3^.

### Turbidostat growth rate measurement

The turbidostat maintained cells growing exponentially between OD_600_ 0.05 and 0.2. In effect, it was run as a batch growth repeater, diluting the culture to OD_600_ 0.05 when it reached OD_600_ 0.2. An exponential line was fit to the growth periods between consecutive dilution events. From the exponential line *I* = *I*_0_ 2^*t/τ*^, the growth rate was determined as *λ* = ln 2*/τ*, where *τ* is the doubling time. The turbidostat spectrometers were blanked with the appropriate medium before each experiment.

### Turbidostat cell size measurement

We fixed cells with a glutaraldehyde and paraformaldehyde mixture and imaged within 24 hr as previously reported^56^, except for the following modifications: 2 *µ*l 25% glutaraldehyde was added to 1 ml 16% paraformaldehyde and cells were resuspended in 300 *µ*l GTE (50mM Glucose 25mM Tris 8.0 10mM EDTA 8.0) per sample after PBS washes.

Before imaging, we adjusted cells to an appropriate cell density as needed. Cells were pipetted onto a 2% agarose pad and briefly dried. The agarose pad was then flipped onto a Willco dish (WillCo Wells, Netherlands) and covered with a glass coverslip to reduce evaporation during imaging. Each experiment consisted of 80-200 images. Sample sizes are presented in Table 5.

We performed fixed cell image analysis with a custom Python script using the OpenCV library. First, we detected contours using an active snakes edge detection algorithm. We then filtered for cell contours using a priori knowledge of cell size and shape, and manually checked for correctly segmented cells. Width and length were calculated from the long and short axis of the cell segments using a simple threshold on the raw phase contrast images. All segmented cells where the width and length fell within 3 standard deviations of the mean for that measurement were kept for further analysis. To calculate cell volume, we assumed the cell was a cylinder with hemispherical ends.

### Turbidostat C period measurement using qPCR

We estimated C period using qPCR and marker frequency analysis. Genomic DNA was prepared from each turbidostat sample using a standard phenol chloroform extraction. We amplified genomic DNA using PowerUp SYBR Green Master Mix (Thermo Fisher Scientific). We used primer pairs targeting chromosomal loci and calculated the C period using the ratio of relative loci copy numbers as discussed previously^3^. Primers are listed in Table 6.

### Mother machine cell preparation and image acquisition

We prepared cultures for mother machine experiments the same as for turbidostat experiments except for the following difference: for translational inhibition experiments, the culture was diluted into the target media with appropriate antibiotics and allowed to grow for several generations before loading into the device.

We performed mother machine experiments as previously described^6, 7^. We used a custom centrifuge to load cells into the growth channels of the mother machine. The time required to remove cells from the water bath shaker, load them into the growth channels, and infuse fresh 37^◦^C media was between 15 and 30 minutes. We then imaged cells for many hours under constant media infusion via a syringe pump (Harvard Apparatus, MA).

For nutrient shift experiments, two syringe pumps were used in conjunction with a manual Y-valve near the device inlet. Cells experienced the change in nutrients in a time interval shorter than the imaging interval^39^.

### Mother machine image processing

Mother machine images were processed with custom Python software. The pipeline takes raw images and produces objects which represent a cell and contains all measured parameters. Briefly, the software aligns and crops images into single channels, segments cells, and links segments in time to construct cell lives and lineages. From the constructed cells we extracted physical parameters in space and time such as size and growth rate. The software has been previously described^7^ with the following modification: segmentation was accomplished with a convolutional neural network implemented with TensorFlow using manually annotated training data^57^.

After segmentation and lineage creation, the resulting cells were filtered for those with measured parameters (septum position, elongation rate, generation time, and birth, division and added length) within 4 standard deviations of their respective population means. We only considered cells in the time interval for which measured parameters and the fluorescent signal were in steady-state. This was normally 3-4 hours after imaging began until imaging ceased. For the growth condition glycerol rich with 3.5 *µ*M chloramphenicol, we excluded cells which divided at the quarter positions, which were less than 5% of all cells. For all conditions, we further selected a subset of cells which could be followed for at least 4-6 consecutive generations. The later filtering step did not affect the parameter distributions, but ensured cell cycle determination was possible in light of the presence of overlapping cell cycles. We only considered mother cells during analysis, but note that other cells along the channel had identical elongation rates.

### Single-cell cell cycle analysis

As described in the main text we used a functional fluorescent DnaN-mGFPmut2 fusion protein. The construct was integrated at the chromosomal locus and expressed under the native promoter. This gene product is the *β*-clamp subunit of DNA polymerase III, which is present at high stoichiometry in active replisomes^54^.

Cell cycle analysis is as described previously^7^. Processed fluorescent images were used to determine the cell cycle parameters manually. We first identified replisome foci in the processed fluorescent images using a Laplacian of Gaussian blob detection method. We then constructed cell traces by plotting cell length versus time, with both the fluorescent signal and foci position projected against the long axis of the cell as demonstrated in Figure **3**. Using an interactive program, we determined the start and end of replication visually based on the position and number of detected foci. For the fastest two growth conditions, glycerol rich with 0 and 1 *µ*M chloramphenicol, termination time and thus C and D period were not determined separately.

### Ensemble cell cycle analysis

In the ensemble method, we aligned cells by size and plotted the ensemble replication state. Based on ours and published measurements, we chose alignment by size as opposed to cell age^11^. To create the ensemble, we find the average number of foci as a function of cell size across all cells. For the slow growing case, the number of foci is 1 at small lengths until a transition period, at which it rises to and plateaus at 2. We take the initiation length to be the length at which the foci count rate of change is the highest, using a differentiation step of 0.2 *µ*m. By inferring the average number of overlapping cell cycles *n*_oc_ from the traces, we can calculate C+D to be C + D = (*n*_oc_ + log_2_(*S*_d_/*S*_i_))⋅τ.

#### Data and code availability

Data and code involved in this study are available upon request.

---

**Extended Figure 1-1:**
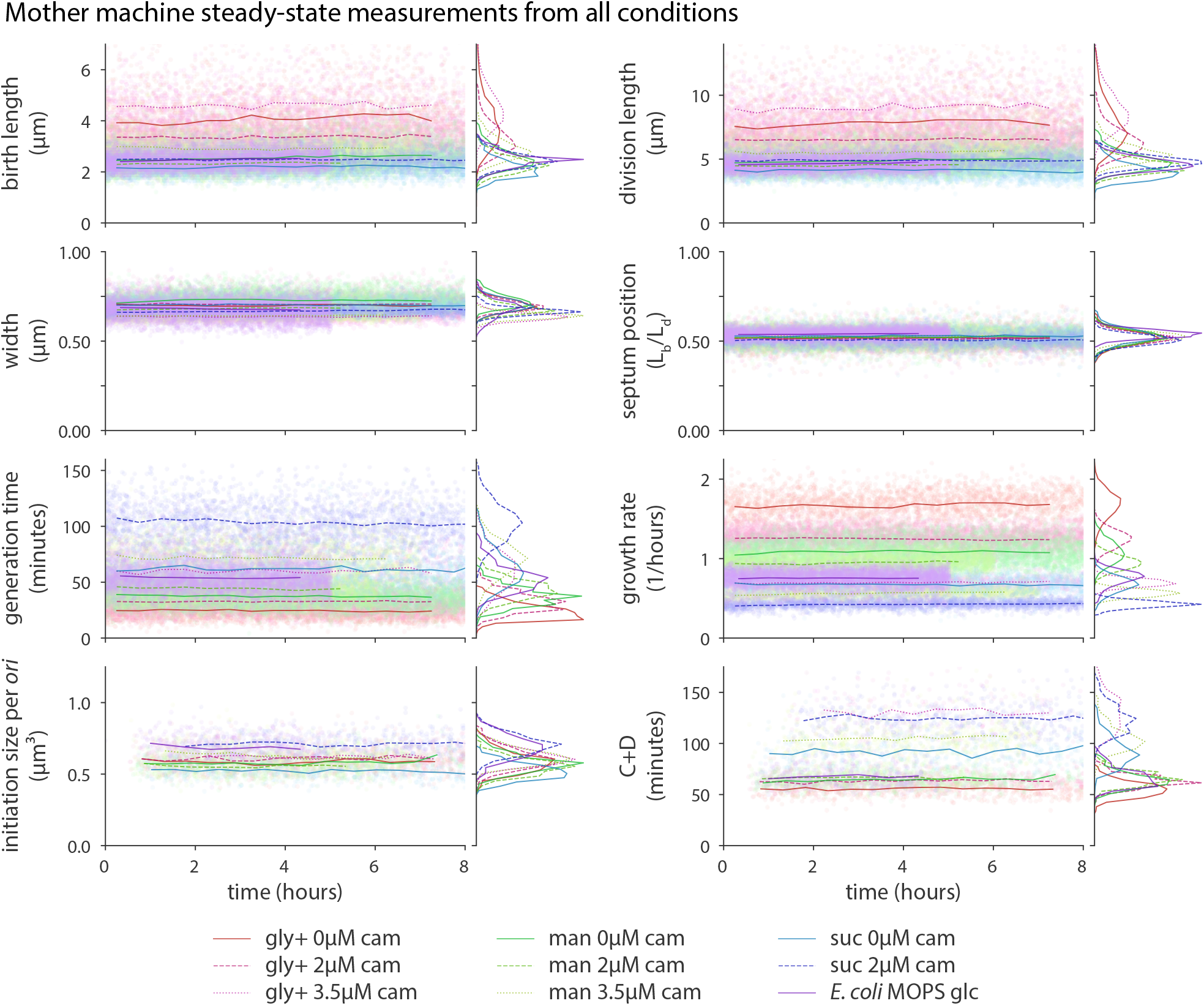
Single-cell steady-state physiological parameters for all conditions. Physiological parameters for all *B. subtilis* mother machine experimental conditions and one *E. coli* experiment. Time course is shown with single-cell measurements (scatter points) and 30 minute binned mean (horizontal lines) plotted against the birth time. Multiple consecutive generations are needed to determine initiation size, C period, and D period, thus a gap exists before those measurements are possible. Single-cell distributions are invariant in time and shown for each condition, sharing the same scale as time course. Colors are as in Figure **1**B. Sample sizes are provided in Table 6.

**Extended Figure 2-1:**
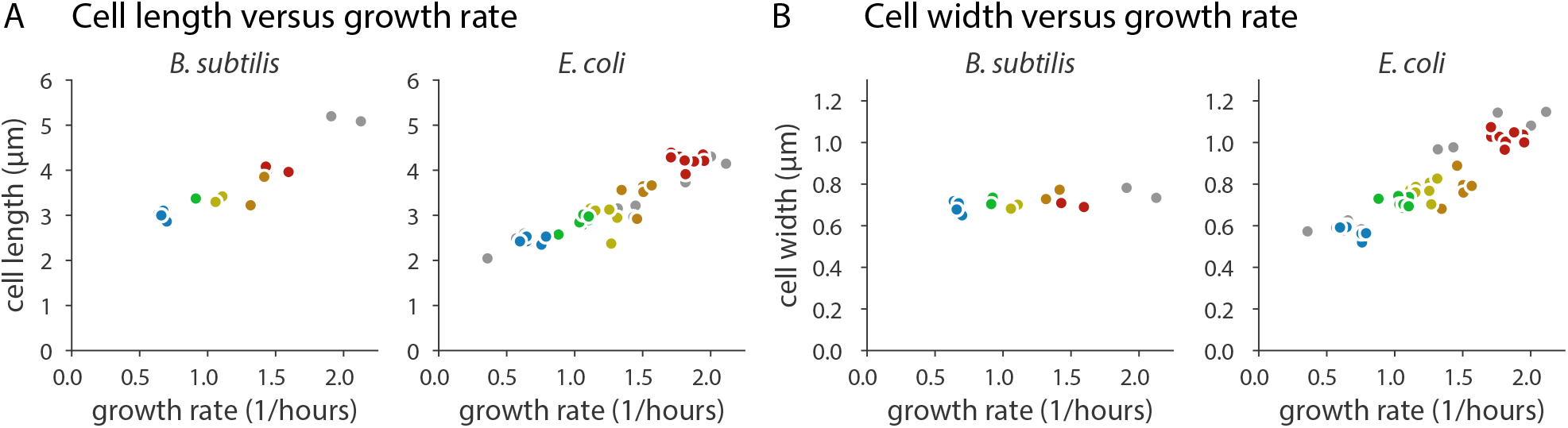
Length and width measurements in *B. subtilis* and *E. coli*. (A) Cell length in *B. subtilis* and *E. coli* increases with growth rate. (B) For *B. subtilis*, width is independent of the nutrient-imposed growth rate. For *E. coli*, width increases with growth rate in a similar manner to length. Colors and conditions are as in Figure **2**. *E. coli* data is from previously published work^3^.

**Extended Figure 2-2:**
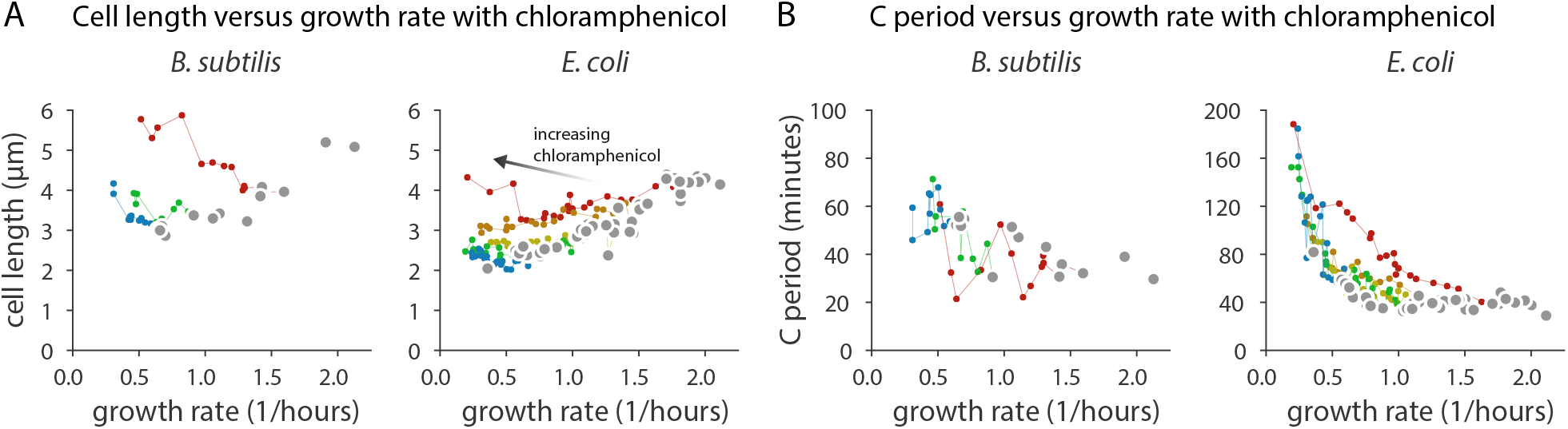
Size and C period under translational inhibition in *B. subtilis* and *E. coli*. (A) Under translation inhibition due to chloramphenicol, the relationship between cell size and growth rate under nutrient limitation breaks down for both *B. subtilis* and *E. coli*. (B) The deviation from the nutrient growth law can be attributed to the change in C period in both species under translation inhibition. Lines connect translation inhibition experiments using the same media. Colors and conditions are as in Figure **2**. *E. coli* data is from previously published work^3^.

**Extended Figure 3-1:**
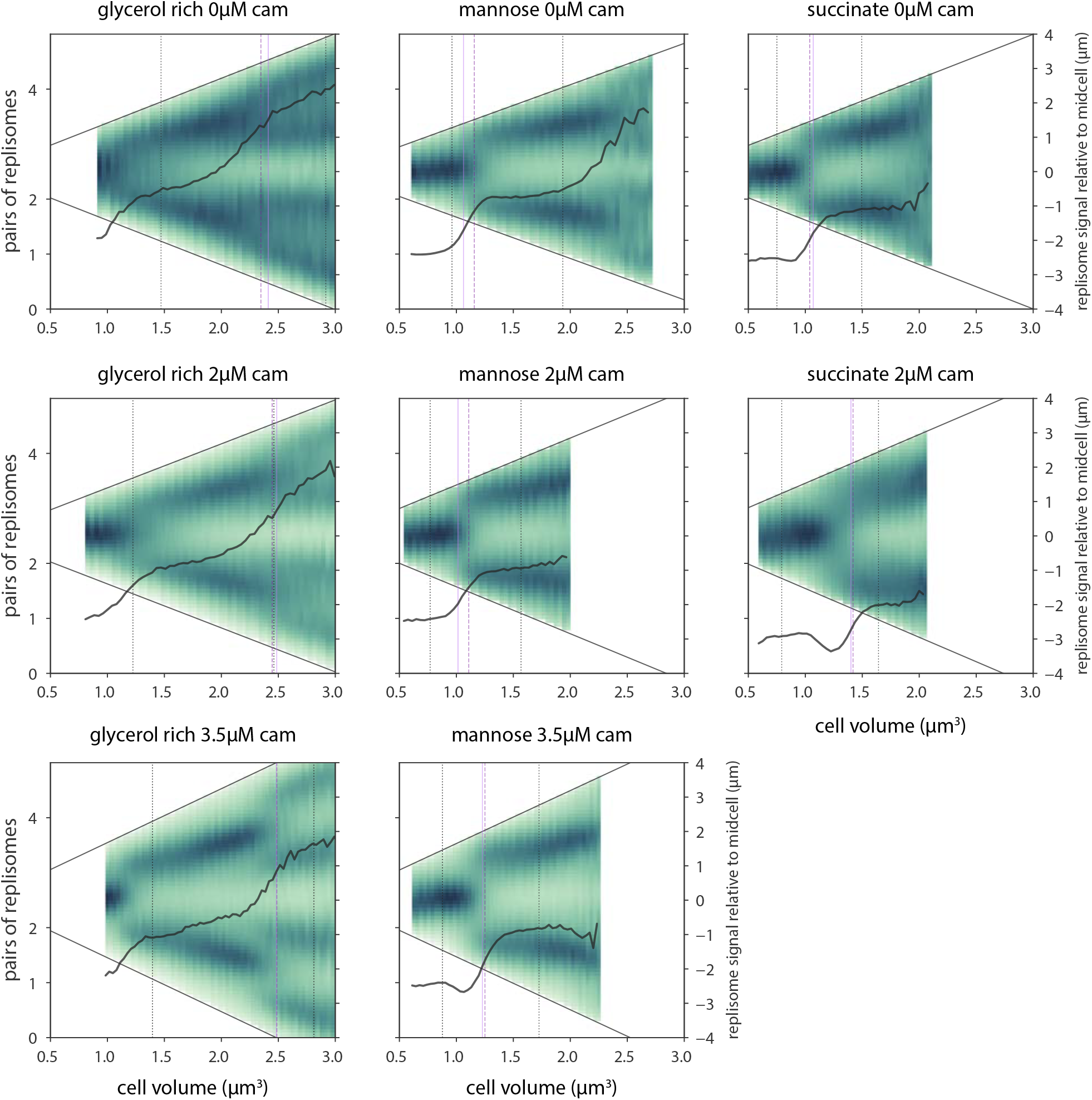
Ensemble replisome count and localization for all conditions. Ensemble plots for three media conditions tested with and without translational inhibition. Average pairs of replisomes (thick black line) is plotted against cell volume with consistent scale across conditions. As in Figure **3**C, purple vertical lines show the initiation size from the average of single cells (dashed) and the ensemble method (solid). Vertical dotted black lines indicate the average birth and division size. We can calculate the average number of foci for sizes outside the average birth and division length due to cell-to-cell variability (ensemble data is shown at sizes to which at least 50 cells contributed). The average number of foci may be above or below the theoretical number. This is because replisomes transiently dissociate, and a pair of replisomes may be counted as two foci when they are not colocalized^24, 58, 59^. The normalized DnaN-mGFPmut2 signal relative to midcell (green background) shows the localization of replisomes over the cell cycle, with the diagonal solid black lines indicating the cell periphery. Termination and replication initiation are often synchronous and correspond to bifurcations in the localization pattern.

**Extended Figure 4-1:**
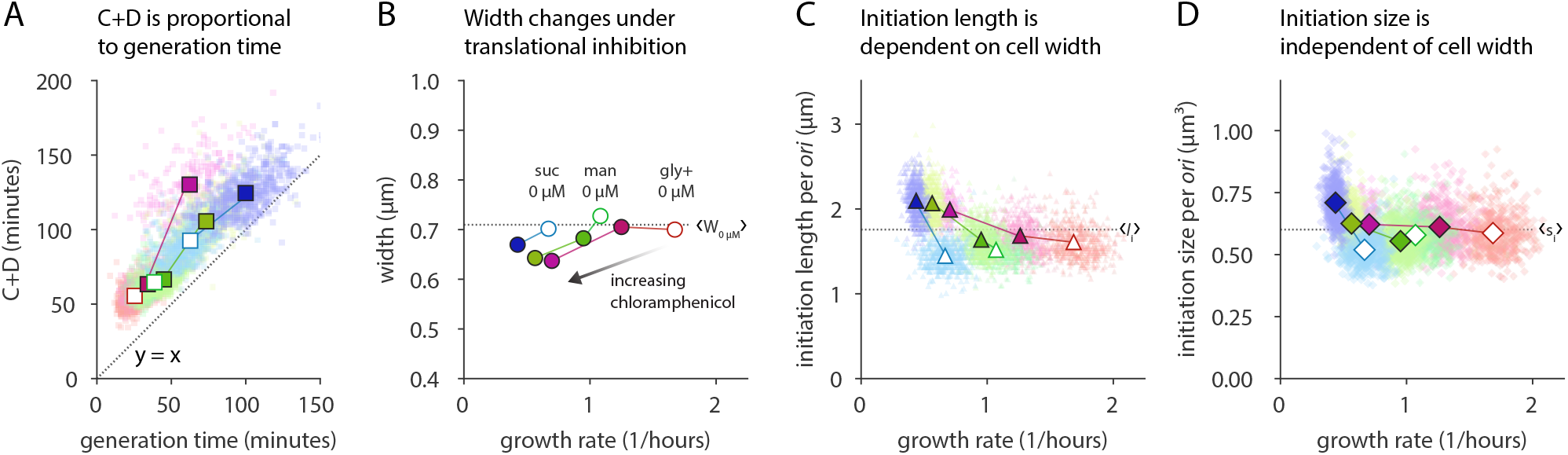
*B. subtilis* cell cycle and initiation size behavior. (A) C+D is proportional to generation time in *B. subtili* when the generation time is modulated by nutrient condition or translational inhibition. (B) Cell width under translational inhibition decreases as compared to the average width without inhibition, <W_0*µ*m_> (dotted black line). (C) Initiation length increase with width under translational inhibition. Mean initiation length per *ori* <L_*i*_> shown as dotted black line. (D) Initiation size from Figure **4**A reproduced for comparison. In all plots, colors are as in Figure **4**, where lines connect the same growth media with and without chloramphenicol. Scatter points are single-cell data and solid symbols are population averages.

**Extended Figure 4-2:**
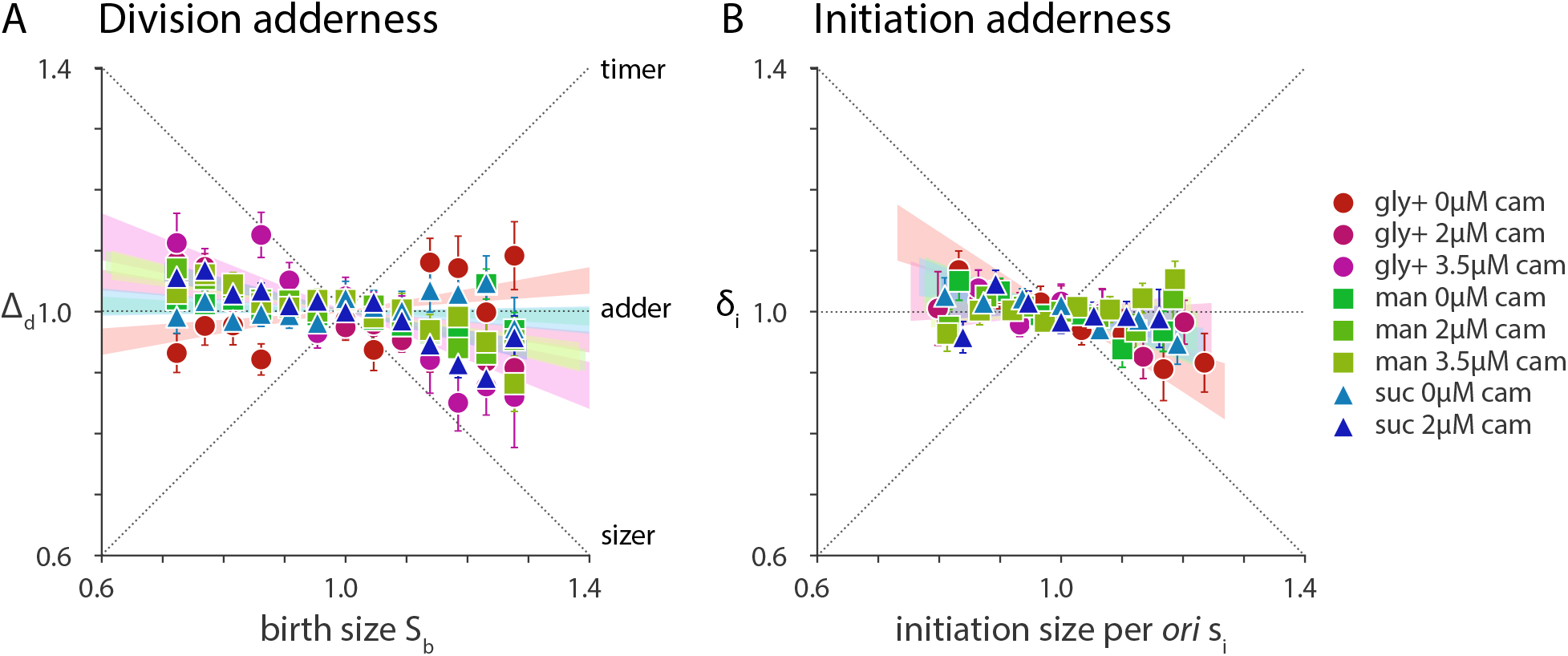
*B. subtilis* is an initiation and division adder. (A) In steady-state growth under nutrient limitation or translational inhibition, *B. subtilis* ∆_d_ is not strongly correlated with the S_b_, consistent with the adder principle. Theoretical correlation between ∆_d_ and S_b_ for the timer, sizer, and adder models of size homeostasis are shown as dotted lines^6^. Symbols are binned data and error bars are standard error of the mean. Shaded wedges are the 95% confidence interval of a linear regression line fit to the underlying data. Data are rescaled by their respective means. (B) *B. subtilis* is an initiation adder such that *δ*_i_ is uncorrelated with s_i_. However, in the fastest growth condition, added initiation size per *ori* behaves sizer-like.

**Extended Figure 5-1:**
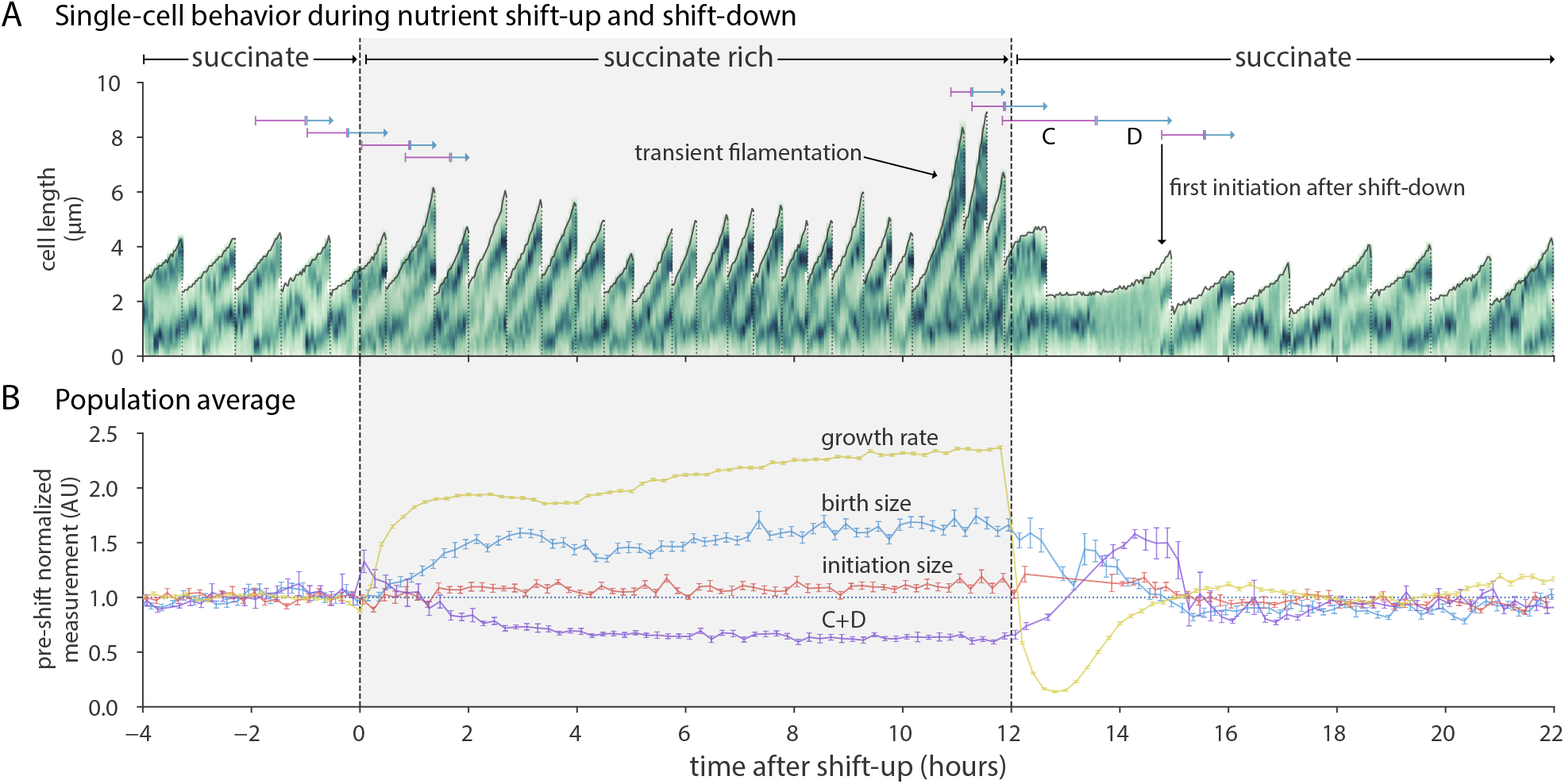
Initiation size is invariant during shift-up and shift-down. (A) Representative lineage trace for cells undergoing shift-up at time zero followed by shift-down 12 hours later. Note that this particular trace exhibits transient filamentation before and unrelated to shift-down. (B) Population average behavior for all cells. Mean lines are calculated as in Figure **5**, except that the measurements are normalized by their respective mean in the 4 hours before shift-up. Additionally, birth size is plotted against the birth time and C+D is plotted against the corresponding division time. Upon shift-up, growth rate immediately increased, simultaneously resulting in an increase in birth size. C+D proportionally decreased. After shift-down, all parameters return to their pre-shift-up average. Despite complex dynamics in these parameters during nutrient shifts, the initiation size showed less than a 10% change during the entire time course. n=7,671 cells (1,695 with initiation size and C+D).

**Extended Figure 6-1:**
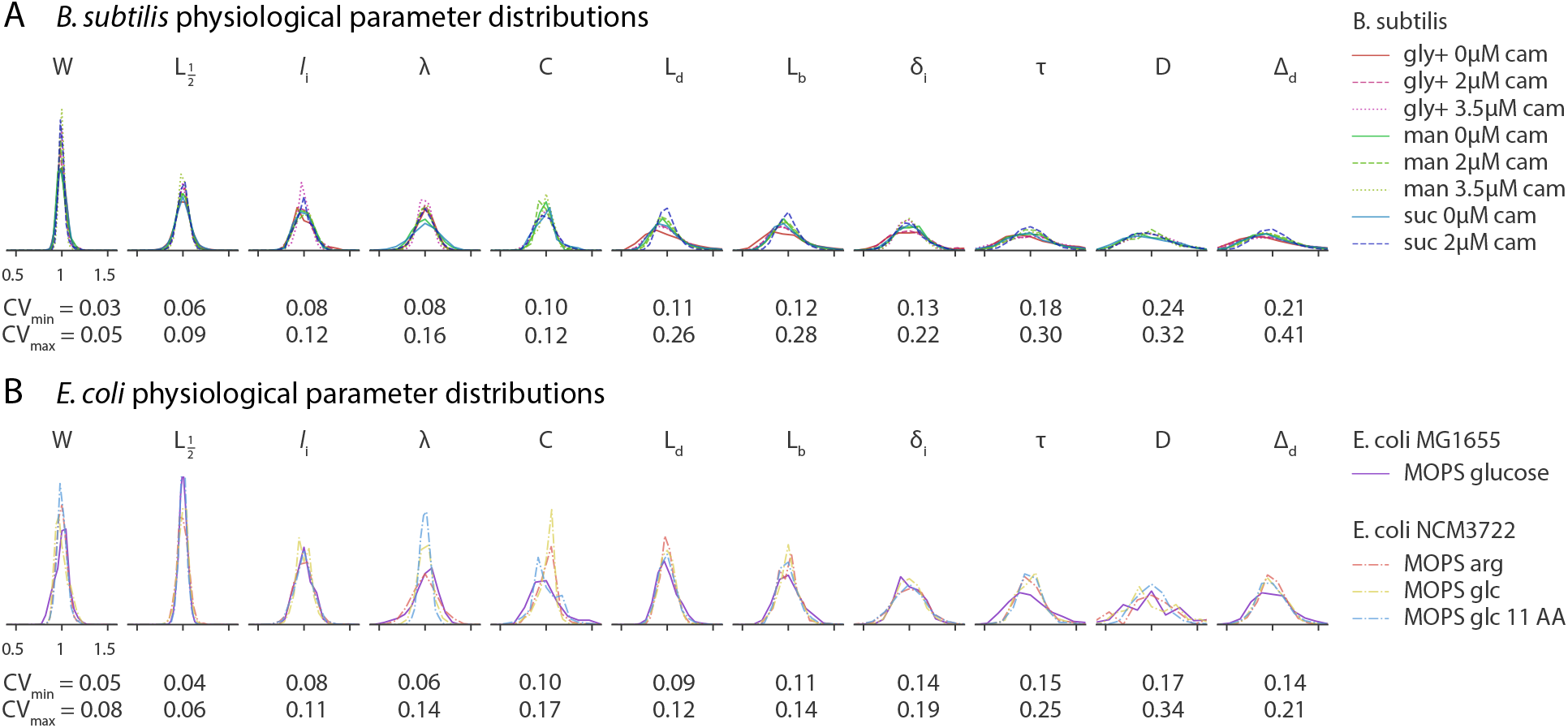
Normalized physiological parameter distributions for all conditions. (A) *B. subtilis* parameter distributions from perturbation experiments are commensurate with nutrient limitation conditions. C period has a smaller CV than D period. (B) In *E. coli*, the CV of C period is smaller than D period. The CV of C+D is smaller than each individually as they are inversely related. Data from *E. coli* NCM3722 grown in MOPS arginine (arg), glucose (glc) and glucose + 11 amino acids (glc 11 AA) are from previously published work^7^.

**Extended Figure 6-2:**
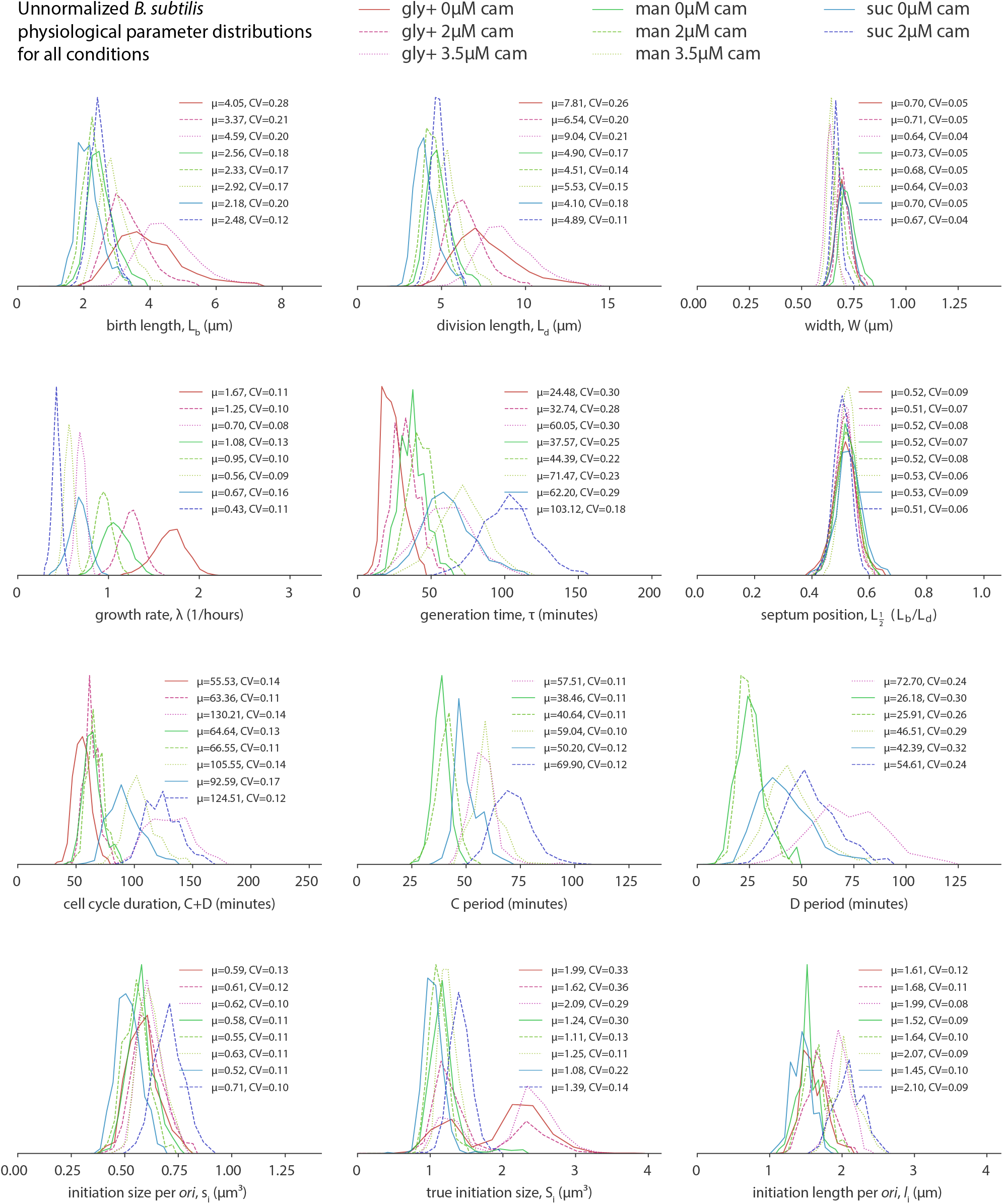
Unnormalized physiological parameter distributions for all conditions. *B. subtilis* physiological parameter distributions for all conditions. Mean (*µ*) and CV are presented in the legends. True initiation size S_i_ is the size at initiation not corrected for the number of *ori*. For conditions in glycerol rich, cells may be born with 1 or 2 replicating chromosomes. Initiation size per *ori* s_i_, width, and septum position are the most conserved across growth conditions.

**Extended Figure 6-3:**
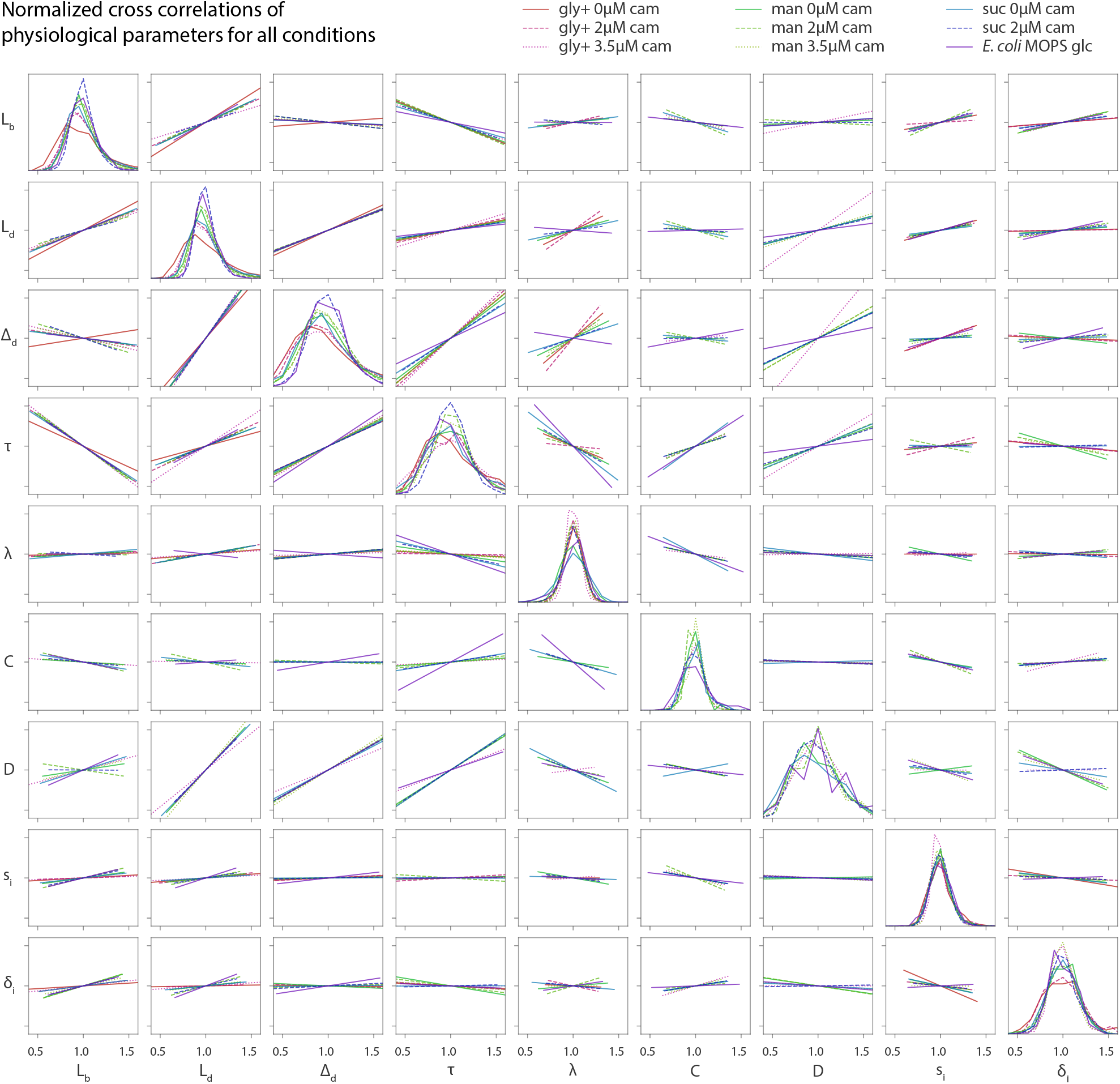
Normalized cross-correlations. Normalized cross-correlations for all *B. subtilis* conditions and one *E. coli* condition. Lines are linear regression fits to the single-cell data. Symbols are as in Figure **6**. Normalized distributions are plotted along the diagonal.

